# Passive Acoustic Monitoring within the Northwest Forest Plan Area: 2023 Annual Report

**DOI:** 10.1101/2025.09.04.674320

**Authors:** Damon B. Lesmeister, Julianna M. A. Jenkins, Natalie M. Rugg, Zachary J. Ruff, Tara Chestnut, Roger Christophersen, Rita Claremont, Raymond J. Davis, Scott Gremel, Aaron Henderson, Edward (Brandon) Henson, Julia Kasper, Heather Lambert, Christopher McCafferty, Steven Mitchell, Sean Mohren, Alex Mueller, Tom Munger, Laura Platt, Dave Press, Courtney Quinn, Suzanne Reffler, Dylan Rhea-Fournier, Madrone Ruggiero, James K. Swingle, Erica Tevini, Alaina D. Thomas, Kirsten Wert

## Abstract

Here we document progress in implementing large-scale passive acoustic monitoring across the Northwest Forest Plan area to track population trends of northern spotted owls (*Strix occidentalis caurina*), barred owls (*S. varia*), marbled murrelets (*Brachyramphus marmoratus*), and a broad array of forest-adapted wildlife. In 2023, we deployed 4,012 autonomous recording units across 1,009 randomly selected 5-km² hexagons, generating nearly 2.2 million hours of recordings, representing approximately 1 petabyte of acoustic data. These data were processed with PNW-Cnet v5, the latest version of our convolutional neural network model, trained on 135 sound classes representing over 80 species and environmental sounds. Model performance demonstrated high precision for focal species and many additional taxa, substantially reducing manual review effort while enabling broad-scale biodiversity assessments. Results confirmed northern spotted owl detections in all 20% sample areas, with occupancy varying geographically and declining notably in the Tyee study area. Barred owls were widely detected, with the highest prevalence in Oregon and Washington and comparatively lower occupancy in California. Marbled murrelets were consistently detected in coastal areas, particularly the Olympic Peninsula and Oregon Coast Range. Beyond these focal species, PAM and PNW-Cnet generated robust datasets for a wide range of birds, mammals, and disturbance indicators, underscoring the value of random-site, multi-species monitoring. The 2023 field season marked the first full implementation of the 2% + 20% NWFP sampling design, expanding monitoring coverage while strengthening collaborations with federal and state partners. These efforts provide the foundation for long-term, cost-effective wildlife monitoring and inform conservation strategies in dynamic forest ecosystems.

## 1. Introduction

Northern spotted owl (*Strix occidentalis caurina*) populations have been monitored since the 1990’s as part of the Northwest Forest Plan (NWFP) Interagency Monitoring Program to assess effectiveness of the plan, and to inform management and conservation decisions. Population monitoring has revealed continued and increasing rates of population decline throughout the northern spotted owl geographic range, as well as identifying barred owls (*S. varia*) and available habitat as important factors associated with those trends (Lesmeister et al. 2018b, Yackulic et al. 2019, Franklin et al. 2021). Two phases were envisioned in the establishment of the monitoring program (Lint et al. 1999). Phase I of northern spotted owl population monitoring would rely on demographic data and Phase II would be based on habitat monitoring if population change were found to follow trends in forests suitable for nesting and roosting (Lint et al. 1999). The study design for Phase I focused on call-back surveys to locate territorial owls on historical study areas comprised primarily of federal lands, then capturing, marking, and resighting those birds to estimate vital rates and population change (Franklin et al. 1996, Lint et al. 1999). In 2020, the NWFP Regional Interagency Executive Committee decided to discontinue Phase I and transition to Phase II over a two-year period. Phase II entails a coupling of habitat monitoring with passive acoustic monitoring survey data to assess trends in the populations (Lesmeister et al. 2021, Lesmeister and Jenkins 2022).

Passive acoustic monitoring using autonomous recording units (ARUs) has been demonstrated to be effective for conducting surveys for northern spotted owl and barred owls (Duchac et al. 2020), distinguishing northern spotted owl sex (Dale et al. 2022), estimating pair status (Appel et al. 2023), integrating with traditional territory survey data (Weldy et al. 2023), and detecting trends in northern spotted owl populations (Lesmeister et al. 2021). Further, ARUs allow for extended-duration sessions, which greatly decreases technician effort in the field while increasing the quantity of data collected (Tegeler et al. 2012). Development of artificial intelligence models to automate detections results in rapid and effective data processing and analysis workflows for northern spotted owl and a wide range of other vocal wildlife species (Ruff et al. 2020, Ruff et al. 2021, Ruff et al. 2023).

Here we provide the 2023 annual progress report on passive acoustic monitoring conducted throughout the northern spotted owl range. We report ARU survey effort in randomly selected 5-km^2^ hexagons, automated detections of 135 sound classes, and validated results for northern spotted owls, barred owls, and marbled murrelets (*Brachyramphus marmoratus*). Details on previous years’ results are available in earlier annual reports (Lesmeister et al. 2018a, Lesmeister et al. 2019, Lesmeister et al. 2022, Lesmeister et al. 2023)

## 2. Study Area

We collected data within 15 national forests, 5 Bureau of Land Management districts, and 9 National Park lands that were primarily under federal ownership and administered by US Forest Service, US Bureau of Land Management, or National Park Service (Fig. 1). Within the federal lands, we collected data within 10 historical northern spotted owl demographic study areas (Franklin et al. 2021) which were the 20% sampling areas. Nine of the study areas (OLY = Olympic Peninsula, CLE = Cle Elum, RAI = Mount Rainier National Park, COA = Oregon Coast Range, HJA = H.J. Andrews Experimental Forest, TYE = Tyee, KLA = Klamath, CAS = Oregon South Cascades, NWC = Northwest California) were long-term demographic study areas for northern spotted owl monitoring under the Northwest Forest Plan (Franklin et al. 2021), one study area (MAR = Marin County) was included due to long-term and ongoing northern spotted owl demographic monitoring (Fig. 1). The remaining federal lands within the NWFP area were sampled as 2% sampling areas, with additional sampling occurring in some locations (WA 2% = Washington 2% sampled areas; OR 2% = Oregon 2% sampled areas; CA 2% = California 2% sampled areas). In 2023, we collected data from 43 designated Wilderness Areas (Table 1).

**Figure 1.**
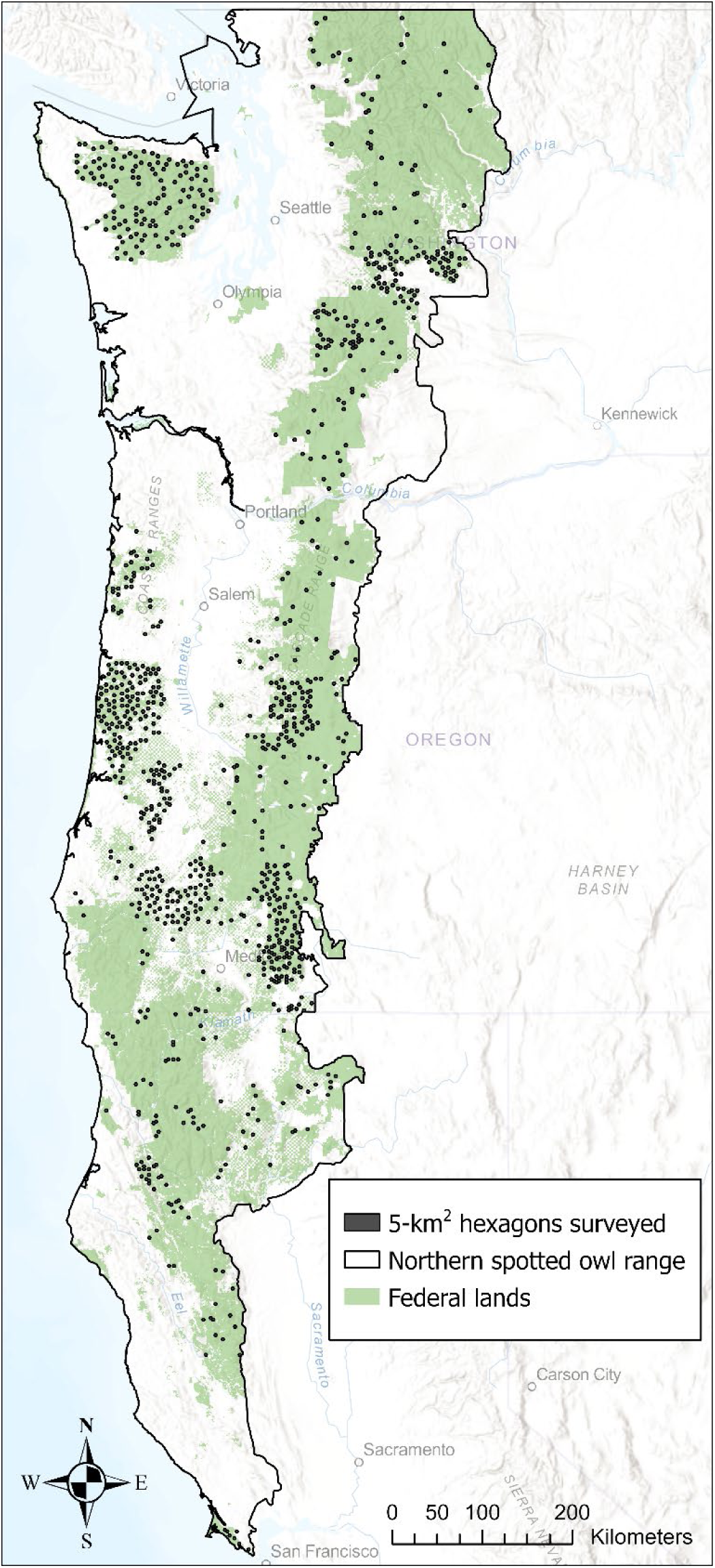
Locations of 5-km^2^ hexagons (*n* =1,009) surveyed on federal lands using passive acoustic monitoring in 2023.

**Table 1.** The number of autonomous recording unit stations surveyed during 2023 in 43 designated Wilderness Areas administered by US Forest Service or US National Park Service.

| Wilderness Area | Stations surveyed |
| --- | --- |
| *US Forest Service* |  |
| Alpine Lakes | 27 |
| Boulder Creek | 1 |
| Buckhorn | 9 |
| Bull of the Woods | 3 |
| Clackamas | 2 |
| Clearwater | 2 |
| Colonel Bob | 8 |
| Cummins Creek | 3 |
| Devil's Staircase | 20 |
| Diamond Peak | 8 |
| Drift Creek | 2 |
| Glacier Peak | 8 |
| Goat Rocks | 8 |
| Indian Heaven | 2 |
| Lake Chelan-Sawtooth | 4 |
| Mark O. Hatfield | 4 |
| Menagerie | 4 |
| Mount Hood | 9 |
| Mount Jefferson | 17 |
| Mount Skokomish | 4 |
| Mount Washington | 2 |
| Mountain Lakes | 10 |
| Noisy-Diobsud | 4 |
| North Fork | 1 |
| Opal Creek | 10 |
| Pasayten | 4 |
| Red Buttes | 4 |
| Rock Creek | 1 |
| Rogue-Umpqua Divide | 10 |
| Salmon-Huckleberry | 1 |
| Siskiyou | 1 |
| Sky Lakes | 63 |
| Snow Mountain | 4 |
| The Brothers | 4 |
| Three Sisters | 49 |
| Trinity Alps | 15 |
| William O. Douglas | 23 |
| Yolla Bolly-Middle Eel | 8 |
| Yuki | 4 |
| *US National Park Service* |  |
| Mount Rainier | 124 |
| Olympic | 238 |
| Phillip Burton | 7 |
| Stephen Mather | 22 |
| TOTAL | 754 |

## 3. Methods

### Sampling design

We created a uniform layer of 5-km^2^ hexagons that covered the entire range of the northern spotted owl (Lesmeister et al. 2021) that is available for download (USFWS 2021). This hexagon size is approximately the size of a northern spotted owl territory core area (Glenn et al. 2004, Schilling et al. 2013) and approximates the home range size reported for barred owls in the Pacific Northwest (Hamer et al. 2007, Singleton et al. 2010, Wiens et al. 2014). Within the historical demographic study areas, we randomly selected 20% of available hexagons that contained ≥50% forest capable lands and ≥25% federal ownership. Outside the historical demographic study areas, we randomly selected 2% of hexagons throughout the entire NWFP area following the same criteria for forest capable lands and federal ownership, stratified by physiographic province. Forest capable lands were those areas with suitable soil type, plant association, and elevation capable of developing into forest (Davis and Lint 2005). In a subset of our study areas (OLY, COA, KLA), we surveyed non-adjacent hexagons to reduce the probability of detecting the same individual in multiple hexagons. Within each hexagon, we deployed 4 ARUs as our sampling stations (Fig. 2).

**Figure 2.**
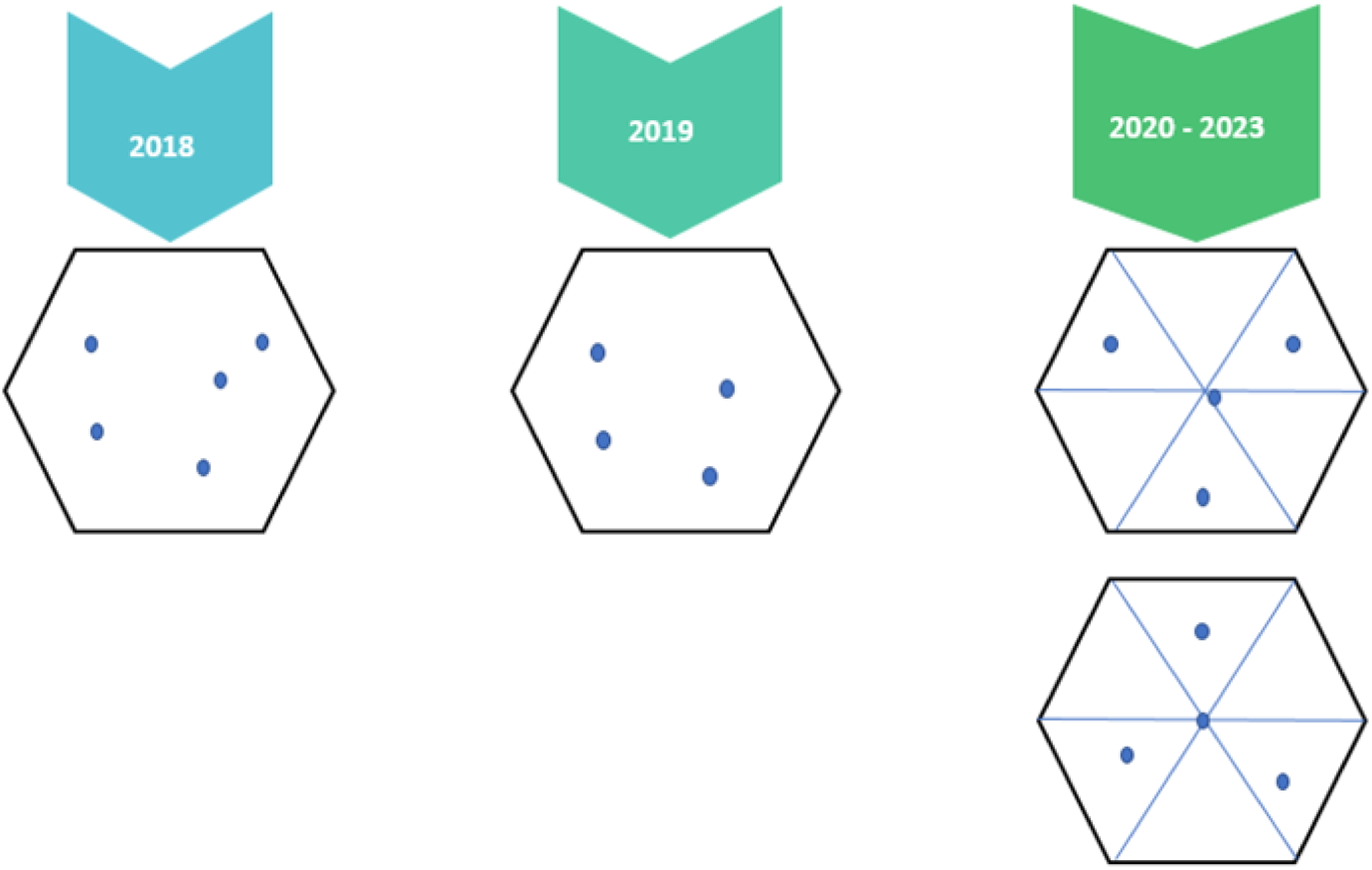
Example sampling station layouts by year. In 2018, five sampling stations were randomly placed within hexagons (no further than 1.5 km from a road or trail) following this rule set: on federal land; mid-to-upper slope positions; ≥ 50 m from roads, trails, and streams; spaced ≥ 500 m apart; and located ≥ 200 m from edge of hexagon. Starting in 2019, established hexagons on COA and OLY had one sampling station randomly removed based on sampling design change, leaving four sampling stations. Newly established hexagons in KLA during 2019 had four random sampling stations selected following within-hexagon placement rule set established in 2018. Newly established hexagons in 2020 - 2023 followed a more standard sampling station layout with one station centrally located and three stations in non-adjacent triangles within the hexagons. Other within-hexagon placement rules established in 2018 was also applied, thus some stations needed to be adjusted to meet rule set requirements.

We collected acoustic data using Song Meter SM4 (primary device with >95% data collected) and Song Meter Mini (Wildlife Acoustics, Maynard, MA) ARUs that are portable, weatherproof, and easily programmable. The SM4s had two built-in omni-directional microphones with signal-to-noise ratio of 80 decibels (dB) typical at 1 kilohertz (kHz), two SDHC/SDXC flash card slots, average of 543 h of recording battery life, and a recording bandwidth of 0.02–48 kHz at levels of -33.5–122 dB. The Song Meter Mini recorded at the same bandwidth, signal-to-noise ratio of 78 dB, one omni-directional microphone, one SDHC/SDXC flash card slot, and 210–1040 h battery life depending on configuration. These ARU models recorded sound with equivalent sensitivity to normal range of human hearing, and their effective listening radius may be affected by external factors such as terrain, vegetation, and weather events such as wind and rain. At each sampling station within a hexagon, we mounted ARUs to small trees (15–20 cm diameter at breast height) to allow microphones to extend past the bole for unobstructed recording ability. We deployed ARUs on federal land; mid-to-upper slope positions; ≥50 m from roads, trails, and streams to reduce vandalism and excessive noise; spaced ≥500 m apart; and located ≥200 m from edge of the hexagon. We programed ARUs to record from 1 h before sunset to 3 h after sunset, 2 h before sunrise to 2 h after sunrise, and for the first 10 min of every hour throughout the day and night (Fig. 3). The deployment of ARUs and data retrieval were the first two steps in our general workflow for collection, processing, and reporting findings (Fig. 4).

**Figure 3.**
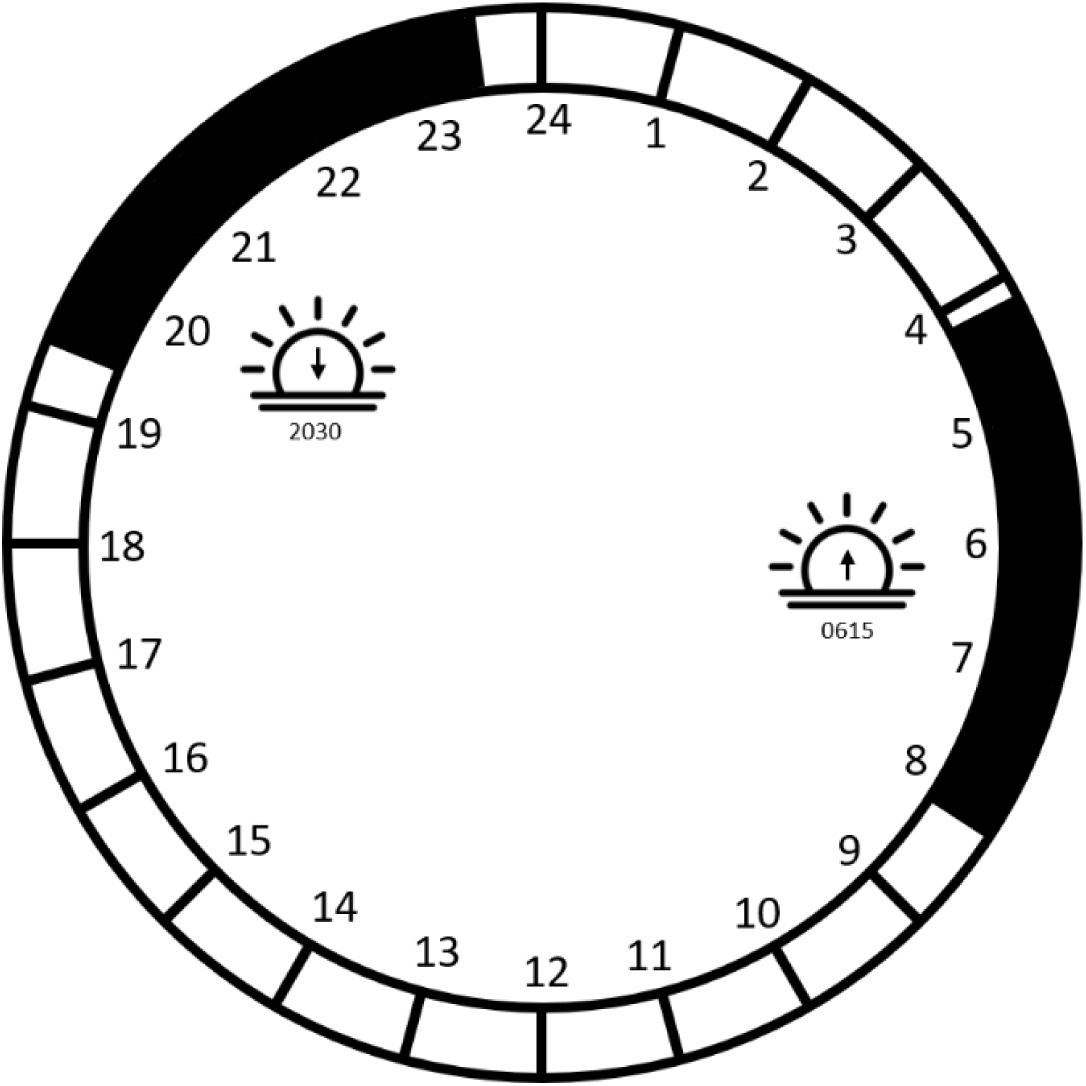
Example 24 h diel cycle (sunrise at 0615, sunset at 2030) recording schedule used on autonomous recording units to conduct passive acoustic monitoring within the Northwest Forest Plan area. Recording times shown with black bars occurring during 4 h blocks during crepuscular period and 10 minutes each hour. The first daily crepuscular block recording starts 2 h before (0415) and ends 2 h after (0815) sunrise, and the second block recording starts 1 h before (1930) and ends 3 h after (2330) sunset.

**Figure 4.**
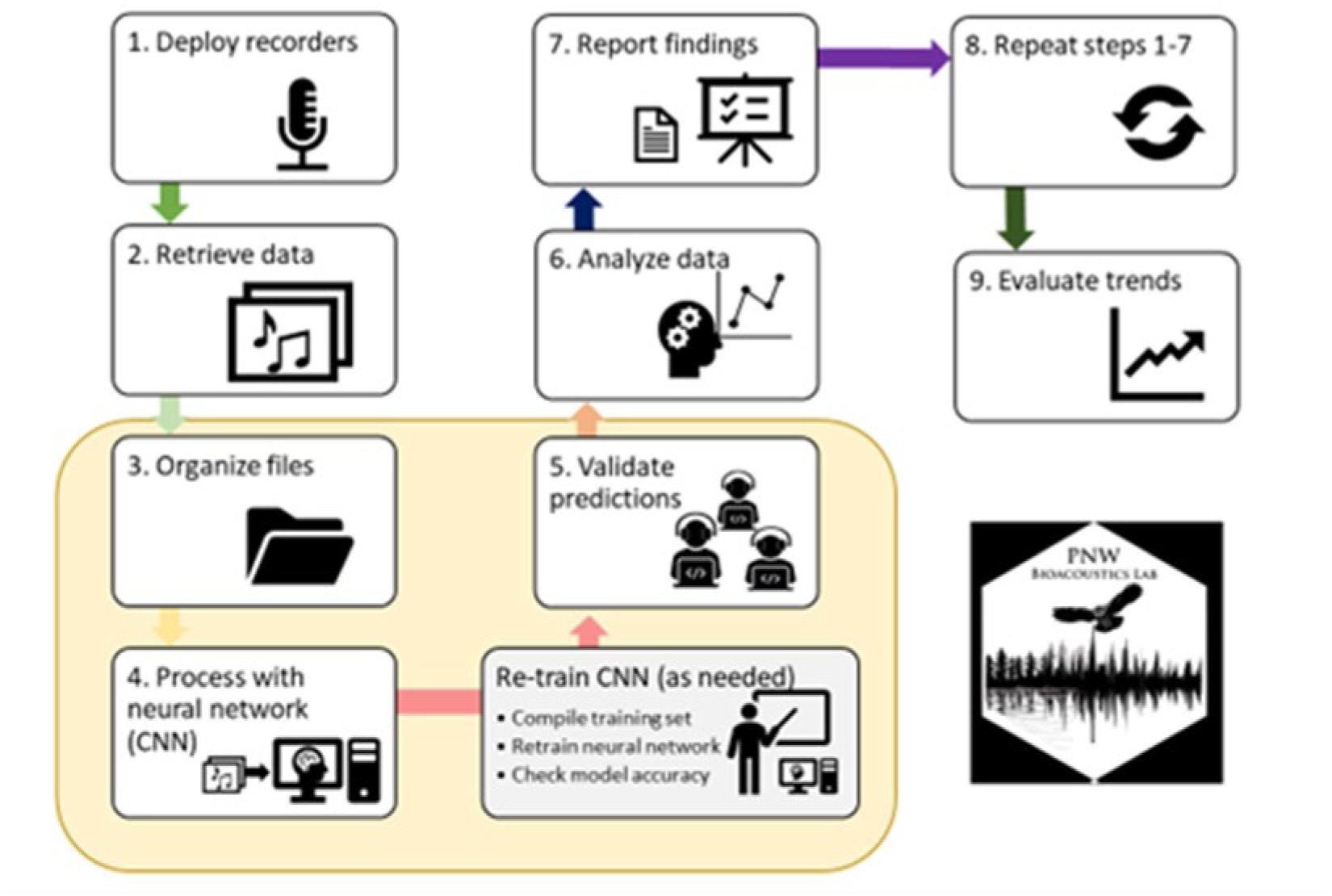
Workflow for the passive acoustic monitoring program within the Northwest Forest Plan area. The process includes data collection, training the convolutional neural network (PNW-Cnet) for automated species identification, processing data, analyzing data, and reporting findings. Highlighted are steps 3–5 which are steps focused primarily on data processing.

### Convolutional neural network (PNW-Cnet) development

Since 2019 we have developed five versions of a convolutional neural network model to automate detections of vocal wildlife species, with each version attaining improved performance and greater number of species identified compared to each preceding version (Fig. 5). Preceding our broadscale passive acoustic monitoring, Duchac et al. (2020) and Duchac et al. (2021) conducted passive acoustic monitoring in 2017 to estimate detection probabilities and occupancy for several owl species in or near three of our study areas (OLY, COA, KLA). From those survey data, Ruff et al. (2020) used the sound clips for six owl species (including northern spotted owl and barred owl) to train our first version of the convolutional neural network (PNW-Cnet v1) to automate species identifications. We replicated and refined the model and process used to train PNW-Cnet v1 for successive PNW-Cnet versions that have been used for data processing.

**Figure 5.**
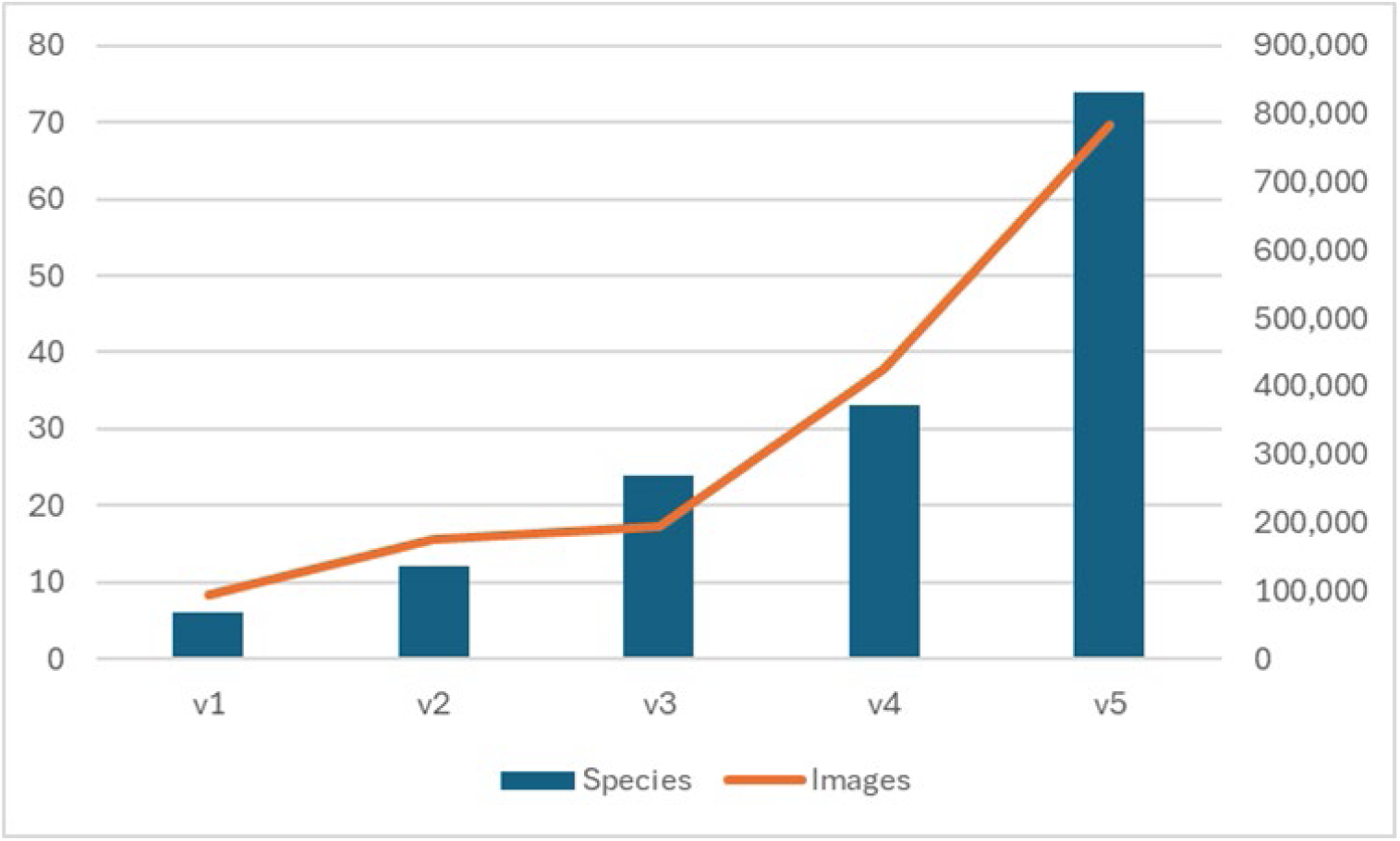
PNW-Cnet version with number of wild species identified to species level (primary y-axis) and sample size of images used in model training set (secondary y-axis) (Ruff et al. 2020, Ruff et al. 2021, Ruff et al. 2023).

Details on development and performance of PNW-Cnet v1 can be found in Ruff et al. (2020). Briefly, we located target species vocalizations in the 2017 data using the Simple Clustering feature of Kaleidoscope Pro software (version 5.0, Wildlife Acoustics) to generate training data. Given that convolutional neural network models are designed for image classification, we split all sound files (.wav) into 12 s segments and then converted those to spectrograms, which are image representations of sound (Fig. 6). We used a 12 s interval because it cleanly divides an hour-long field recording and is long enough to fully contain any of the owl calls. To reflect the variation found in field recordings, we generated multiple spectrograms with different parameters for each 12 s clip (Ruff et al. 2020).

**Figure 6.**
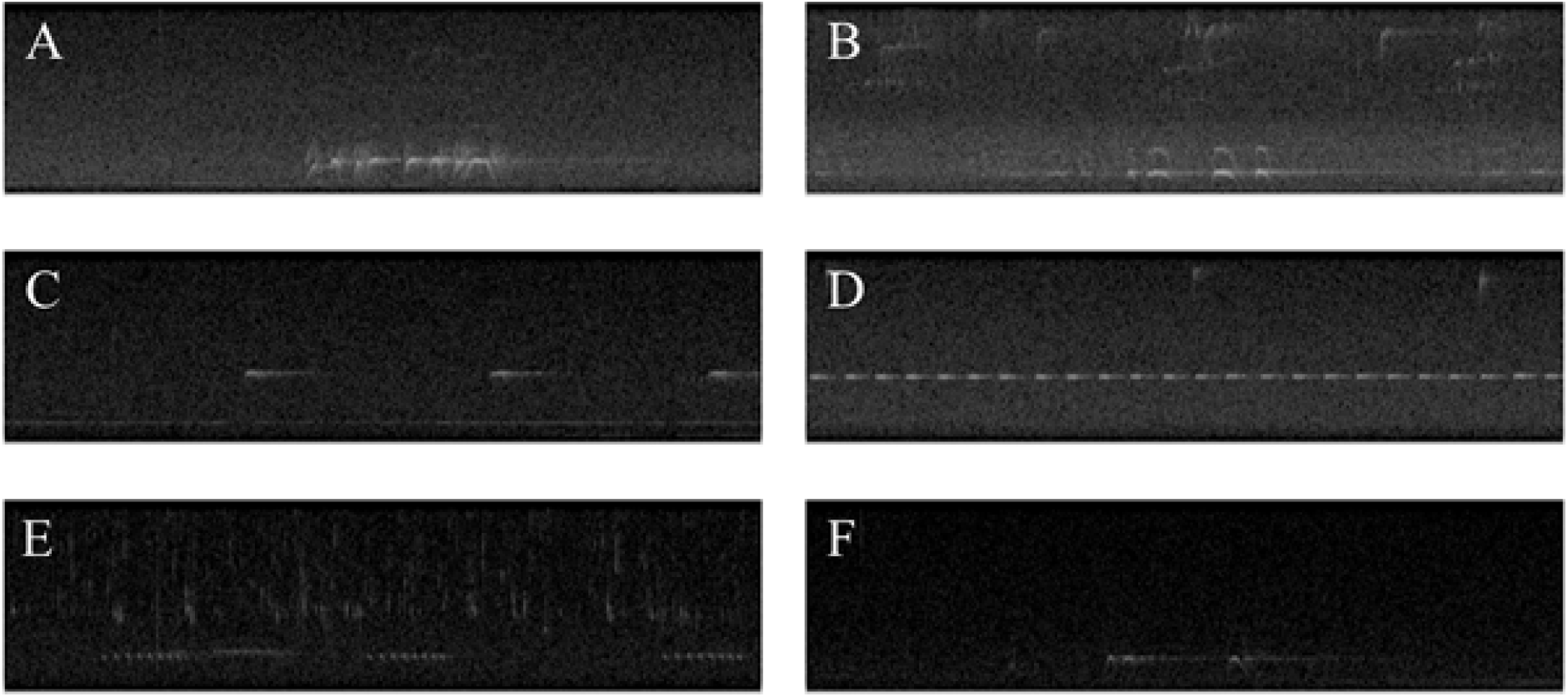
Example spectrogram images of target species calls used by Ruff et al. (2020) to train PNW-Cnet to detect owl calls in field recordings. A = barred owl, B = great horned owl, C = northern pygmy owl, D = northern saw-whet owl, E = western screech owl, F = northern spotted owl. Each spectrogram is 500 x 129 resolution and represents 12 s of audio in the frequency range 0-3000 Hz. Spectrograms like those shown were used in PNW-Cnet v1 and v2. From PNW-Cnet v3 and v4, spectrograms were 1000 x 257 resolution and included the frequency range 0-4000 Hz. Lighter areas represent greater sound intensity.

The final training dataset used by Ruff et al. (2020) included spectrograms for seven target classes: northern saw-whet owl (*Aegolius acadicus*; *n* = 10,003), great horned owl (*Bubo virginianus*; *n* = 9,999), northern pygmy-owl (*Glaucidium californicum*; *n* = 10,003), western screech-owl (*Megascops kennicottii*; *n* = 10,004), northern spotted owl (*n* = 22,373), barred owl (*n* = 22,204), and noise (*n* = 10,003). We implemented the PNW-Cnet v1 model in Python using Keras, an application programming interface, to Google’s TensorFlow software library (Abadi et al. 2015). We trained the PNW-Cnet v1 for 100 epochs on our 94,589 training images, of which 80% were used for training and 20% were set aside as a validation set. See Ruff et al. (2020) for details on model performance for each species.

We found that PNW-Cnet model performance could be improved for target species by increasing the size of the model training dataset and incorporating additional target classes, including vocalizations easily confused with the existing classes. From 2018–2023, we expanded and further developed the PNW-Cnet for processing data with important advancements and expansions occurring each year of the study (Fig. 5). PNW-Cnet v5 includes 135 sound classes, of which are 80 different species, and was trained on 782,918 training images, with about 90% used for training and 10% for validation.

To determine the performance of each version of the PNW-Cnet to correctly classify each sound class, we calculated precision and recall, generated from a test set of clips that were fully tagged by human technicians. Precision is the rate of true positives among apparent detections (clips with an output prediction ≥0.95). Recall is the proportion of calls in the dataset that were detected and correctly identified. To measure class-specific performance for PNW-Cnet v5, we set aside 5% of the intended training dataset (n = 41,202 images), which were fully labeled, and classified these images using the model immediately following training. For each class, we tallied: 1) the number of true positive detections, i.e., clips that contained the class and were assigned a score ≥0.95 for that class; 2) the number of apparent detections, i.e., all clips that were assigned a score ≥0.95 for that class; and 3) the number of available examples, i.e., all clips that contained the class. We calculated precision as true positives divided by apparent detections and recall as true positives divided by available examples.

### Data processing

We followed a multi-step workflow that integrated the latest version of PNW-Cnet to efficiently process large volumes of audio data, combining automated identification and human validation (Ruff et al. 2021). This workflow reduced the necessary human effort by >99% compared to full manual review of the data while producing detection/non-detection data based only on human-confirmed detections for chosen classes. PNW-Cnet generates likelihoods (interpretable as probabilities between 0–1) for each sound class for each 12 s clip. We report the number of estimated detections (i.e., number of clips with score exceeding a score threshold) for each target sound class by study area adjusted by model class precision when available.

Generally, precision estimates at the 0.95 threshold were much higher than the recall estimates. As you reduce the review score threshold, recall scores generally increase and precision declines. For example, precision and recall for northern spotted owl four-note calls using the 0.5 threshold vs 0.95 threshold is 0.87 and 0.80, and 0.98 and 0.36, respectively. As we were most concerned with avoiding false positives, we set the threshold high (0.95) to maximize precision over recall for our report summary.

### Data validation and sex predictions

Output from PNW-Cnet v5 (used to process 2023 data) was validated through a process of review by trained human technicians (Fig. 4). The human validation process consisted of reviewing 12 s audio clips that met our model prediction threshold. We used the program Kaleidoscope Pro for validating PNW-Cnet output by examining the audio and spectrogram to confirm or correct the model classified sound class. Depending on species-specific objectives and need to generate training data for future versions of the PNW-Cnet, we reviewed sound classes at one of four intensities:

1. Fully review all clips,
2. Confirm species detection/non-detection for each ARU station during each week of survey,
3. Confirm annual detection/non-detection at each ARU station,
4. Confirm annual detection/non-detection at hexagon level (detection on any ARU station within the hexagon).

Prior to 2023, we fully reviewed all PNW-Cnet output that scored at or above 0.25 probability threshold for northern spotted owl location calls. In 2023, we fully reviewed the northern spotted owl location call class ≥0.25 in approximately 12% of hexagons reviewed, then switched to a probability threshold of ≥0.50 once we were confident that we were not affecting occupancy estimates by reducing the threshold for northern spotted owl. We reviewed marbled murrelet and barred owl call classes at a threshold of ≥0.95 to confirm weekly ARU-level occupancy.

All hexagons with <100 northern spotted owl calls classified by the initial validator were reviewed by a second reviewer for confirmation of northern spotted owl presence. High quality spectrograms with confirmed northern spotted owl calls were analyzed to determine aspects of frequency and call length and these values were used to make out-of-sample predictions from a linear model (Dale et al. 2022) to determine the sex of the vocalizing owl. We classified 95% prediction intervals (PI) <0.5 as females and those with 95% PI > 0.5 as male, and predictions not meeting either of these criteria were classified as unknown sex. Hexagons with > 100 northern spotted owl calls or those with evidence of female or pair vocalizations were reviewed again by expert reviewers. Expert reviewers confirmed model predicted female calls and assigned female presence at sites with counter calling, or non-territorial calls overlapping male 4- note hoots. We report the proportion of surveyed hexagons with validated detections for northern spotted owls, marbled murrelets, and barred owls.

### Removal of data affected by call-back surveys

Call-back surveys for northern spotted owl and barred owl were commonly used in our study areas by biologists working on other research projects (e.g., Franklin et al. 2021, Wiens et al. 2021) and project-level clearance surveys. These surveys broadcast recorded calls of northern spotted owl, or other target species to elicit a territorial response. Beginning in 2021, we distributed a recording consisting of a brief series of pure tones (1 s at 0.5, 1.5 and 1.0 kHz) for call-back surveyors to voluntarily play at the same volume directly before or after northern spotted owl call-back surveys (USFWS 2021). We requested and received broadcast survey information from surveyors in or around our sampling locations at the end of each field season. We manually validated all ≥0.95 predicted detections of the survey tone from PNW-Cnet v5. We removed any validated detections of the surveyed species if there was a reported or suspected call-back survey in the hexagon on the same night, or if we could identify that the detection was a call-back survey auditorily.

### Background noise analysis

Background noise has consistently been found to be an important predictor of detection probabilities in species occurrence models (Duchac et al. 2020, Duchac et al. 2021). Therefore, we used the sound pressure level analysis feature in Kaleidoscope Pro to quantify background noise levels at each ARU sampling station. We created weekly estimates of average daily background noise for each ARU station which are available upon request but not reported here.

## 4. Results

Here we present the most up to date results that should be considered preliminary with need for further validation and quality-assurance and quality-control before formal analyses can be conducted. The amount of data collected and coverage within the NWFP area has increased each year of passive acoustic monitoring since 2018 (Table 2). In 2023, we surveyed a total of 4,012 sampling stations in 1,009 hexagons (Table 2) with nearly 2.2 million h of recordings (Table 3) and approximately 1 Petabyte of data collected. We conducted surveys in five national parks (Crater Lake, Olympic, North Cascades, Mount Rainier, and Redwood), five Bureau of Land Management districts in Oregon, and 14 national forests (Deschutes, Fremont-Winema, Gifford Pinchot, Klamath, Six Rivers, Mendocino, Mount Baker-Snoqualmie, Mt. Hood, Okanogan-Wenatchee, Olympic, Rogue River-Siskiyou, Shasta-Trinity, Siuslaw, Umpqua, and Willamette). Deployments took place from February 24 – July 27. In 2023, 754 of our sampling stations were in designated Wilderness Areas administered by US National Park Service (n = 391) or US Forest Service (n = 363; Table 1). Each year we experience some degree of data loss by various sources such as wildfire, theft, animal damage, and software or hardware failure. In 2023, barred owl, northern spotted owl and great gray owl (*S. nebulosa*) call-back surveys were reported within or adjacent to our sampled hexagons, and any detections of those species overlapping the night of the audio survey were removed.

**Table 2.** Passive acoustic monitoring effort during 2018–2023 within in the Northwest Forest Plan Area, summarized by historical study area (i.e., 20% sampling) and 2% sampling (outside 20% sample density) by state. COA = Oregon Coast Range, OLY = Olympic Peninsula, KLA = Klamath, CLE = Cle Elum, TYE = Tyee, HJA = H.J. Andrews Experimental Forest, CAS = Oregon South Cascades, NWC = Northwest California, MAR = Marin County, RAI = Mount Rainier National Park.

| Area | Number of hexagons |  |  |  |  |  | Number of stations |  |  |  |  |  |
| --- | --- | --- | --- | --- | --- | --- | --- | --- | --- | --- | --- | --- |
|  | 2018<sup>a</sup> | 2019 | 2020 | 2021 | 2022 | 2023 | 2018<sup>a</sup> | 2019 | 2020 | 2021 | 2022 | 2023 |
| COA | 120 | 106 | 120 | 120 | 117 | 120 | 578 | 413 | 471 | 475 | 466 | 476 |
| OLY | 88 | 120 | 119 | 119 | 120 | 120 | 436 | 472 | 464 | 472 | 474 | 479 |
| KLA |  | 63 | 73 | 73 | 73 | 73 |  | 244 | 290 | 276 | 292 | 292 |
| CLE |  |  | 69 | 75 | 75 | 75 |  |  | 267 | 298 | 299 | 300 |
| TYE |  |  |  | 40 | 40 | 40 |  |  |  | 155 | 160 | 160 |
| HJA |  |  |  | 70 | 70 | 70 |  |  |  | 279 | 280 | 278 |
| CAS |  |  |  | 98 | 98 | 98 |  |  |  | 380 | 391 | 391 |
| NWC |  |  |  | 30 | 30 | 30 |  |  |  | 120 | 119 | 118 |
| MAR |  |  |  | 7 | 7 | 7 |  |  |  | 26 | 27 | 27 |
| RAI |  |  |  | 11 | 24 | 24 |  |  |  | 43 | 96 | 96 |
| WA 2%<sup>b</sup> |  |  |  |  | 23 | 79 |  |  |  |  | 91 | 312 |
| OR 2%<sup>b</sup> |  |  |  |  | 13 | 166 |  |  |  |  | 52 | 661 |
| CA 2% |  |  |  |  |  | 76 |  |  |  |  |  | 299 |
| TOTAL | 208 | 289 | 381 | 643 | 690 | 1,009 | 1,014 | 1,129 | 1,492 | 2,524 | 2,747 | 4,012 |

<sup>a</sup> During 2018 survey design was five stations per hexagon.
<sup>b</sup> In 2022, Gifford Pinchot National Forest in Washington and Umpqua National Forest in Oregon were sampled at 2%.

**Table 3.** Thousands of hours of passive acoustic monitoring data collected during 2018–2023 in each study area and processed for automated species identification with PNW-Cnet. COA = Oregon Coast Range, OLY = Olympic Peninsula, KLA = Klamath, CLE = Cle Elum, TYE = Tyee, HJA = H.J. Andrews Experimental Forest, CAS = Oregon South Cascades, NWC = Northwest California, MAR = Marin County, RAI = Mount Rainier National Park. WA 2%, OR 2%, and CA 2% were data collected in each state on the 2% sampling outside the 20% sampling density on historical study areas.

| Study area | 2018<sup>a</sup> | 2019 | 2020 | 2021 | 2022 | 2023 |
| --- | --- | --- | --- | --- | --- | --- |
| COA | 197 | 155 | 187 | 217 | 258 | 259 |
| OLY | 147 | 184 | 219 | 215 | 264 | 271 |
| KLA |  | 92 | 130 | 135 | 162 | 171 |
| CLE |  |  | 104 | 153 | 152 | 168 |
| TYE |  |  |  | 78 | 92 | 88 |
| HJA |  |  |  | 131 | 150 | 149 |
| CAS |  |  |  | 178 | 188 | 195 |
| NWC |  |  |  | 48 | 54 | 62 |
| MAR |  |  |  | 11 | 10 | 15 |
| RAI |  |  |  | 11 | 51 | 71 |
| WA 2%<sup>b</sup> |  |  |  |  | 44 | 192 |
| OR 2%<sup>b</sup> |  |  |  |  | 30 | 385 |
| CA 2% |  |  |  |  |  | 151 |
| TOTAL | 344 | 431 | 640 | 1,177 | 1,456 | 2,179 |

<sup>a</sup> During 2018 survey design was five stations per hexagon.
<sup>b</sup> In 2022, Gifford Pinchot National Forest in Washington and Umpqua National Forest in Oregon were sampled at 2%.

### PNW-Cnet v5

PNW-Cnet v5 marked a significant increase in the number of species (n = 80) from the 37 in PNW-Cnet v4 (Ruff et al. 2023) and had high precision for most sound classes (Table 4). Using PNW-Cnet v5, we were able to generate predicted detections of all focal and most nontarget species (Table 4). PNW-Cnet v5 precision was high (>0.91) for all northern spotted owl, barred owl, and marbled murrelet call types (Table 4). Precision was also high for anthrophony sounds and other important management species such as corvids (Table 4). We also observed high performance for most of the mammals, but sample size remains low for many of those species (Table 4). Recall was low for many sound classes, which suggests larger training data are needed to improve model performance for recall. There were 24 PNW-Cnet v5 classes that had no apparent detections (spectrograms assigned scores ≥0.95) in the test set or in the 2023 data (Table 4). As precision is defined as the proportion of true positives among apparent detections, it was not possible to estimate precision for these classes at the 0.95 score threshold. These were all new classes whose performance will likely increase with future versions of PNW-Cnet. There were 108 sound classes with precision estimates >0.90 at the 0.95 threshold (recall ranging from 0.01–0.97).

**Table 4.**
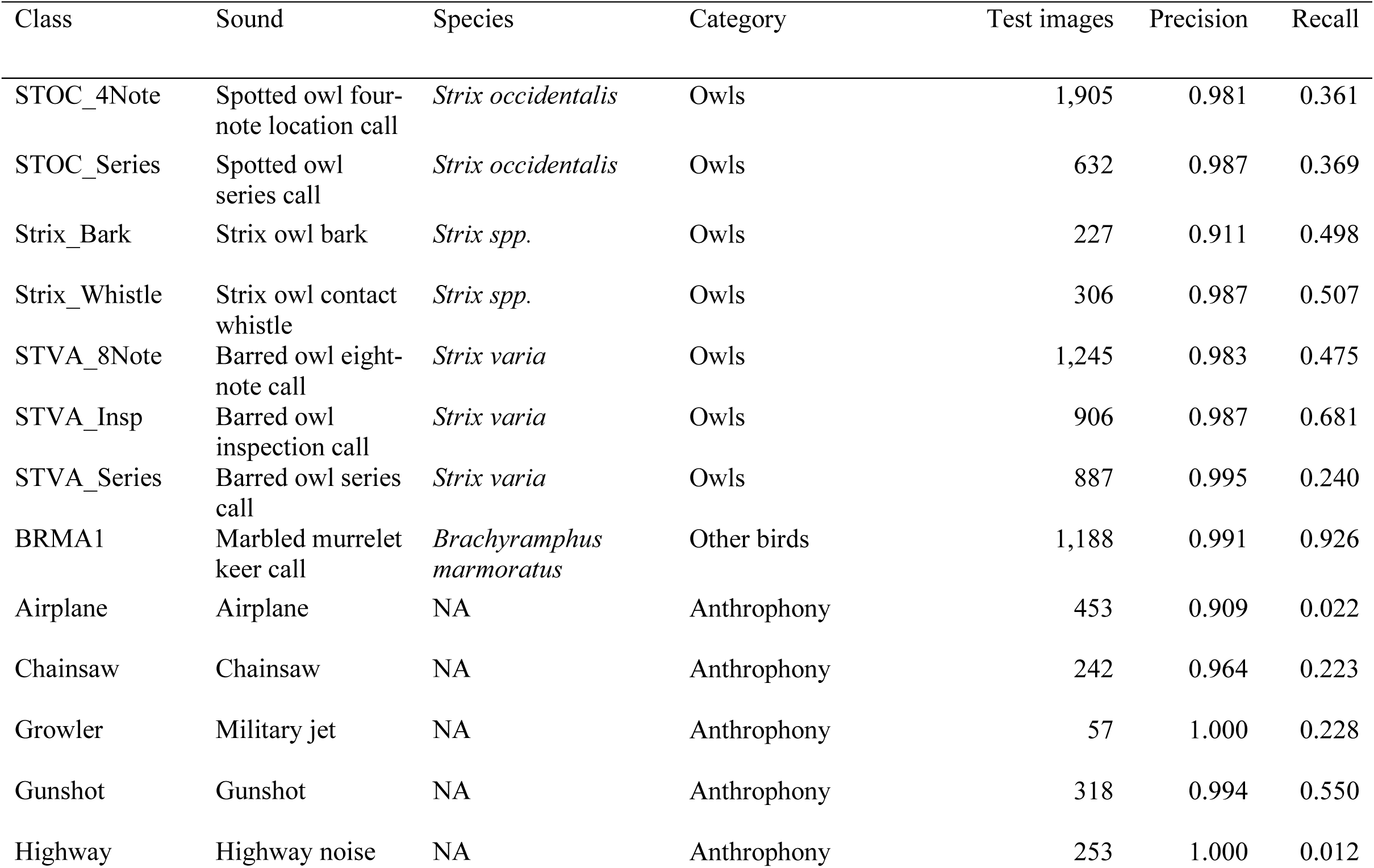
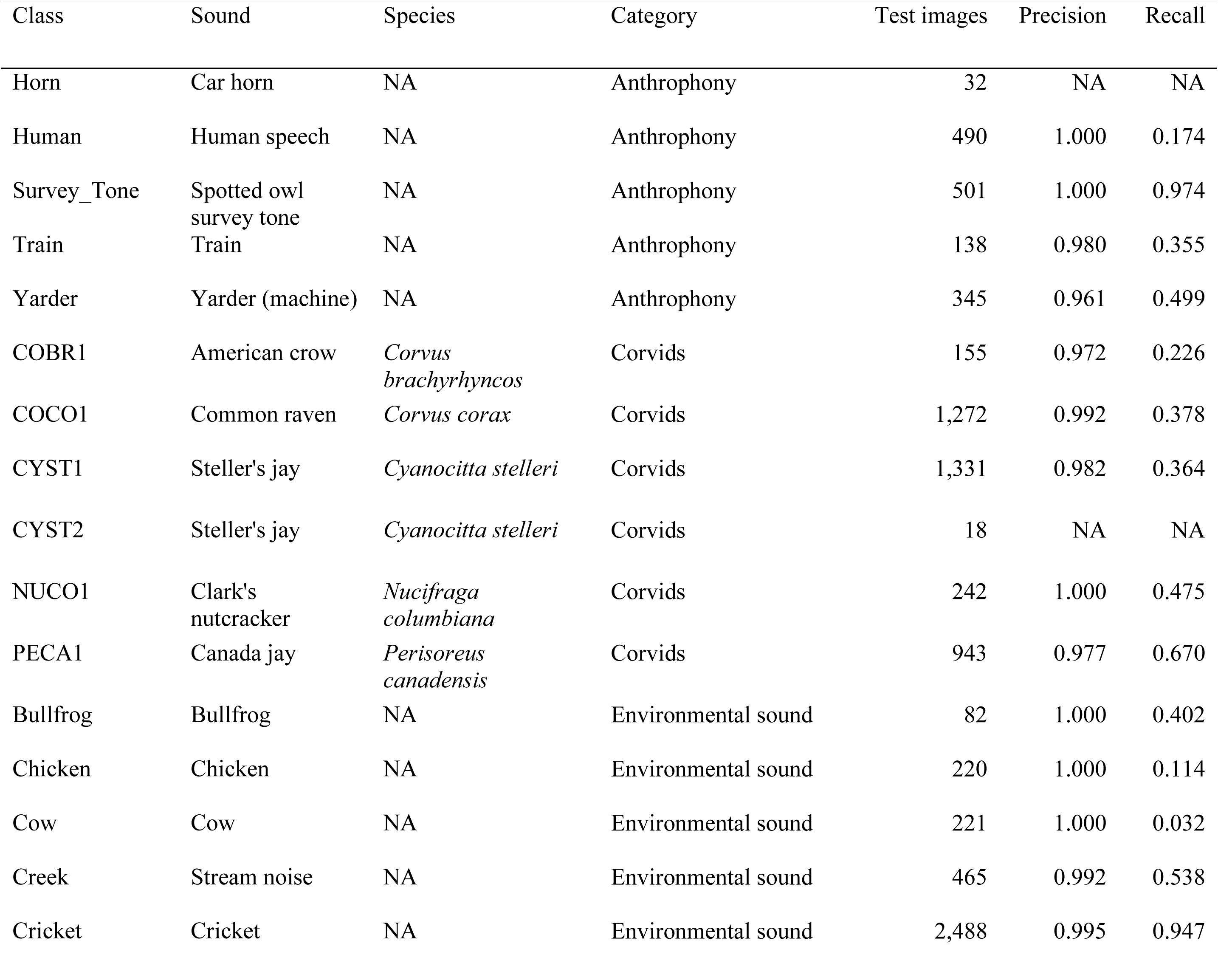
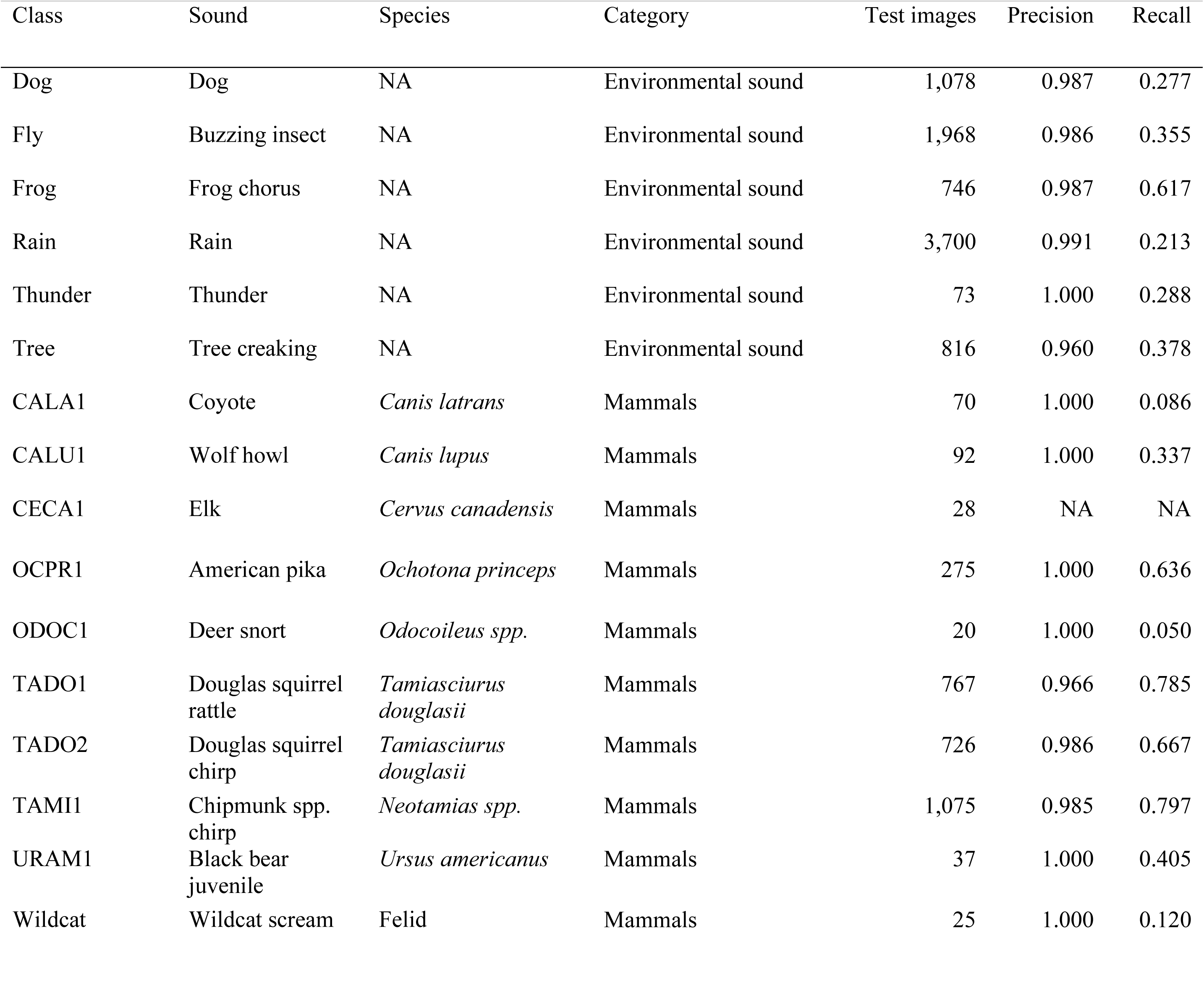
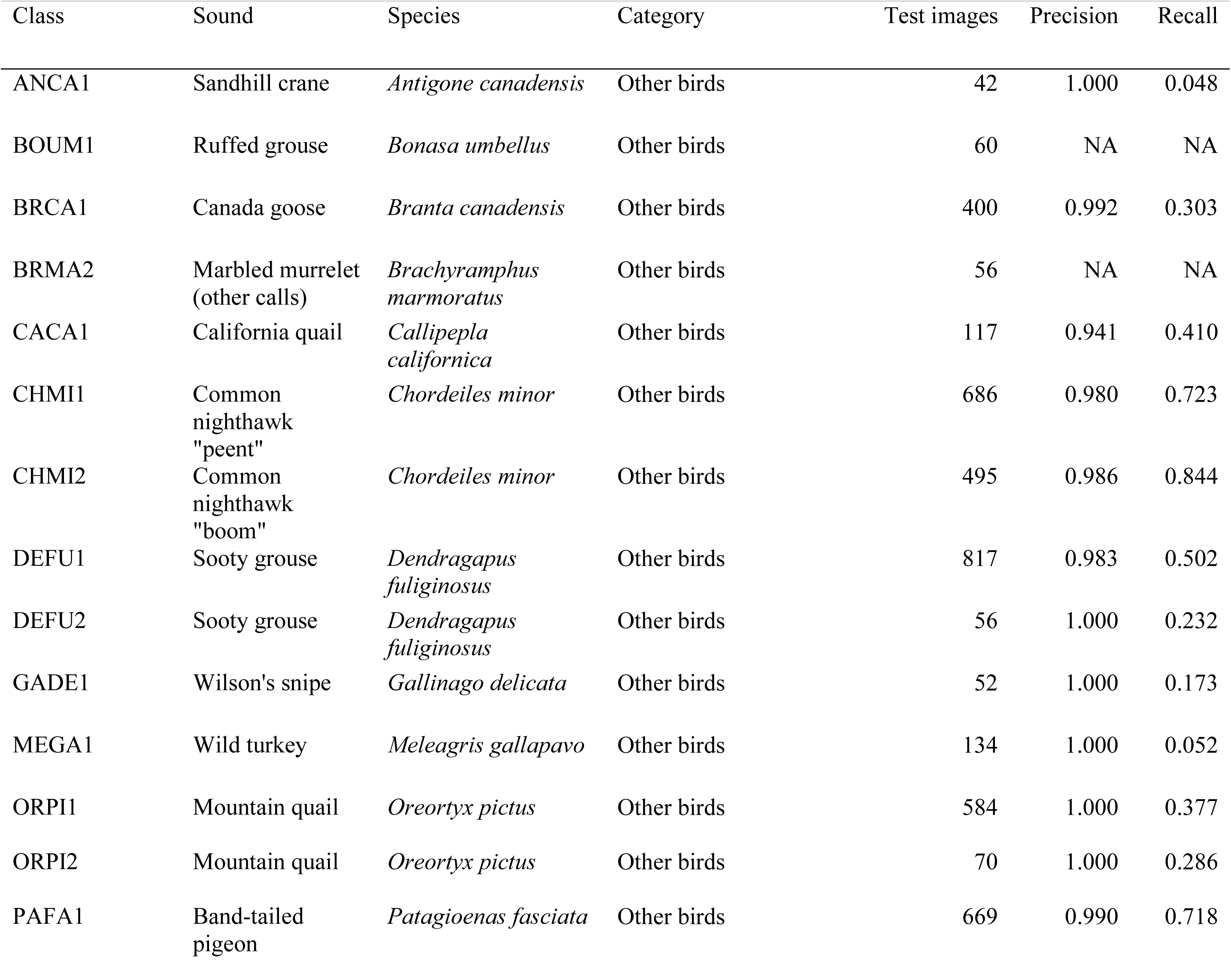
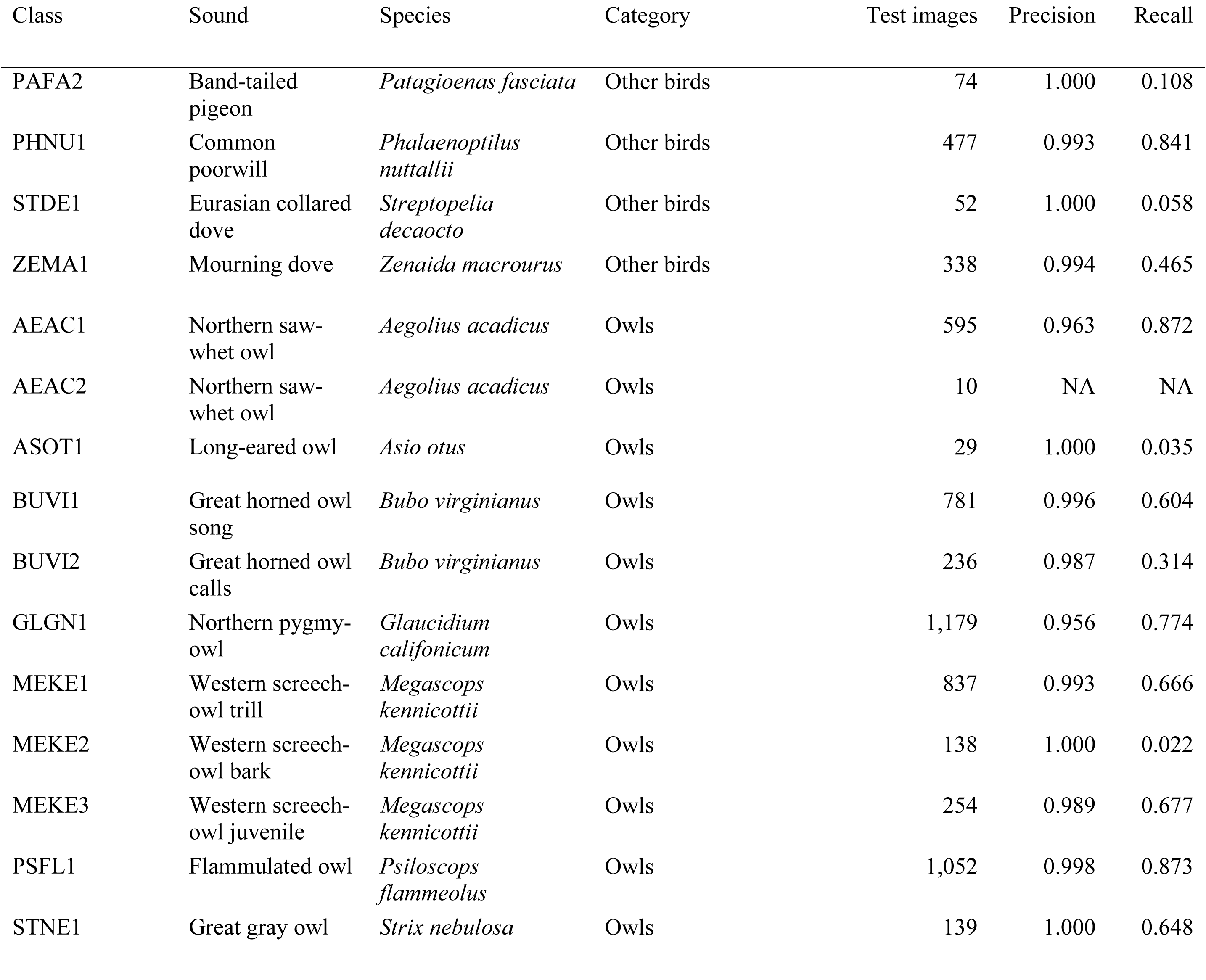
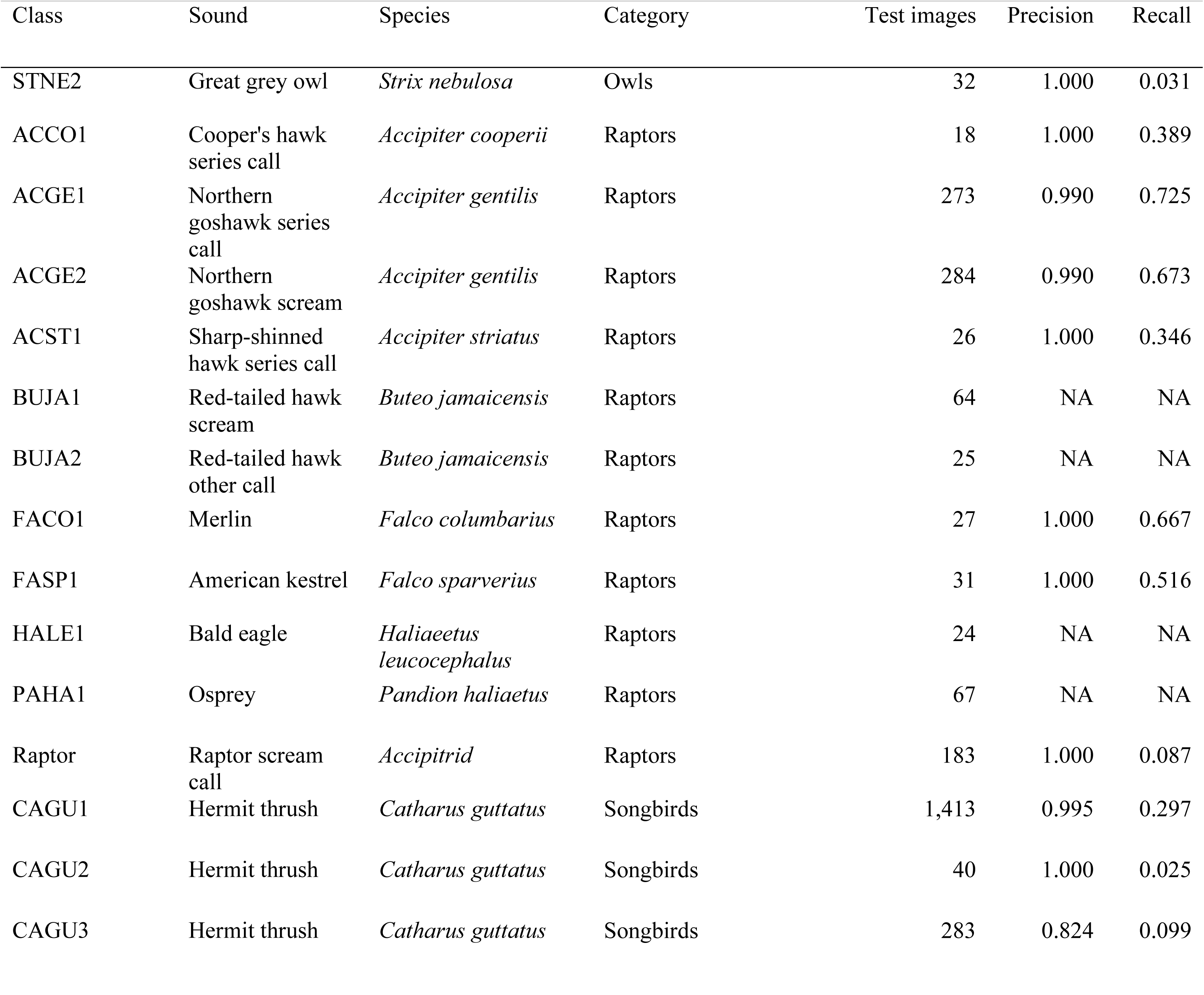
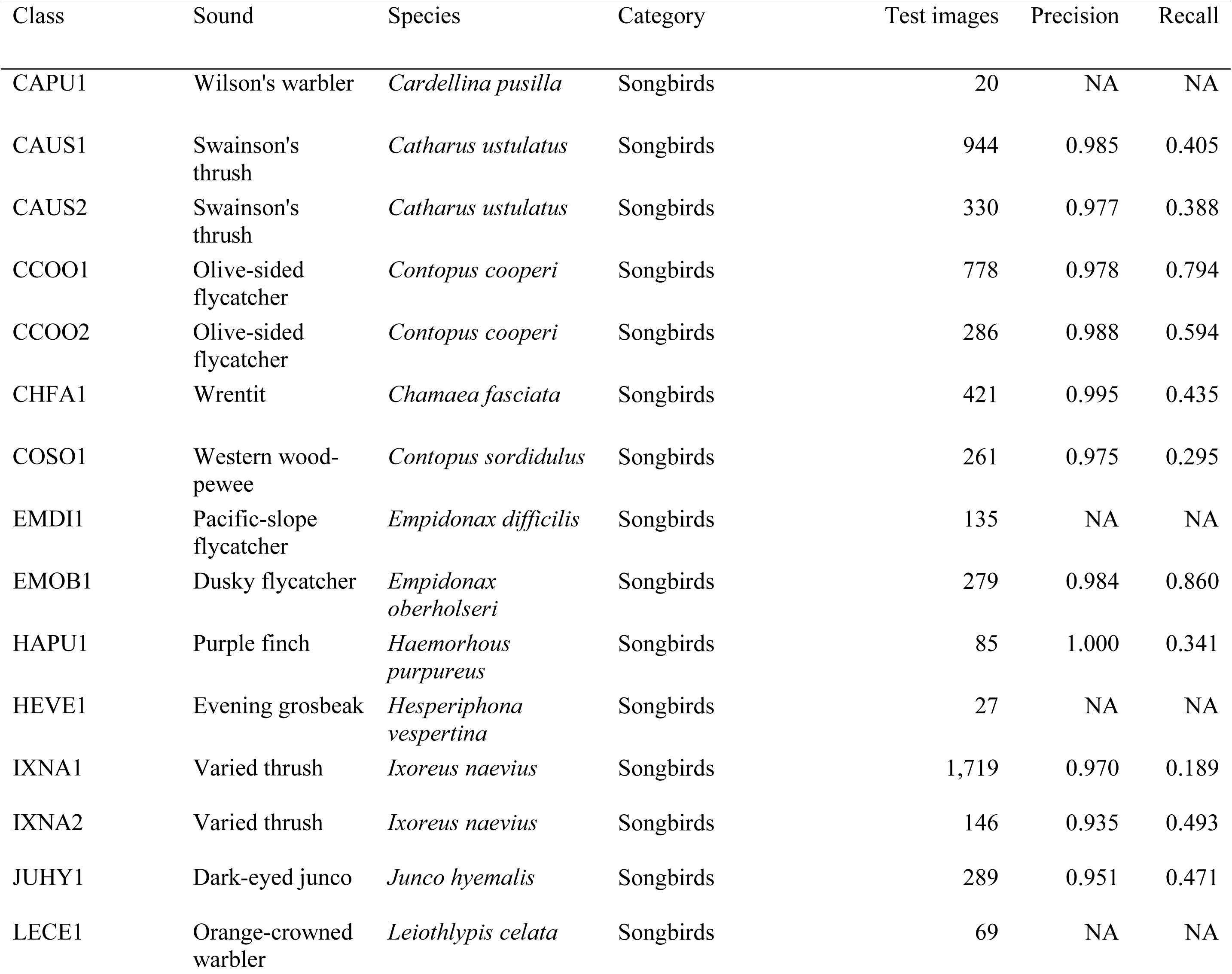
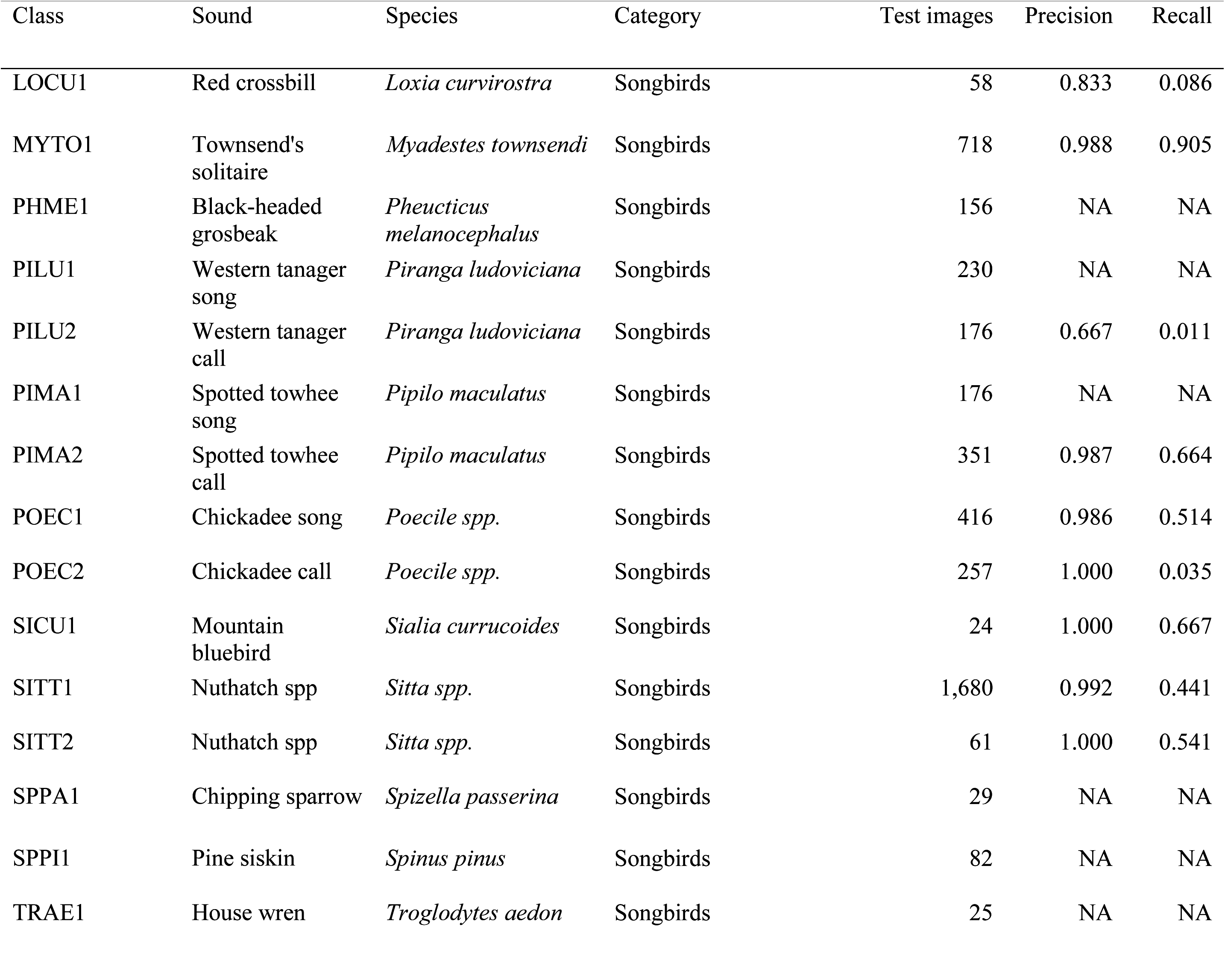
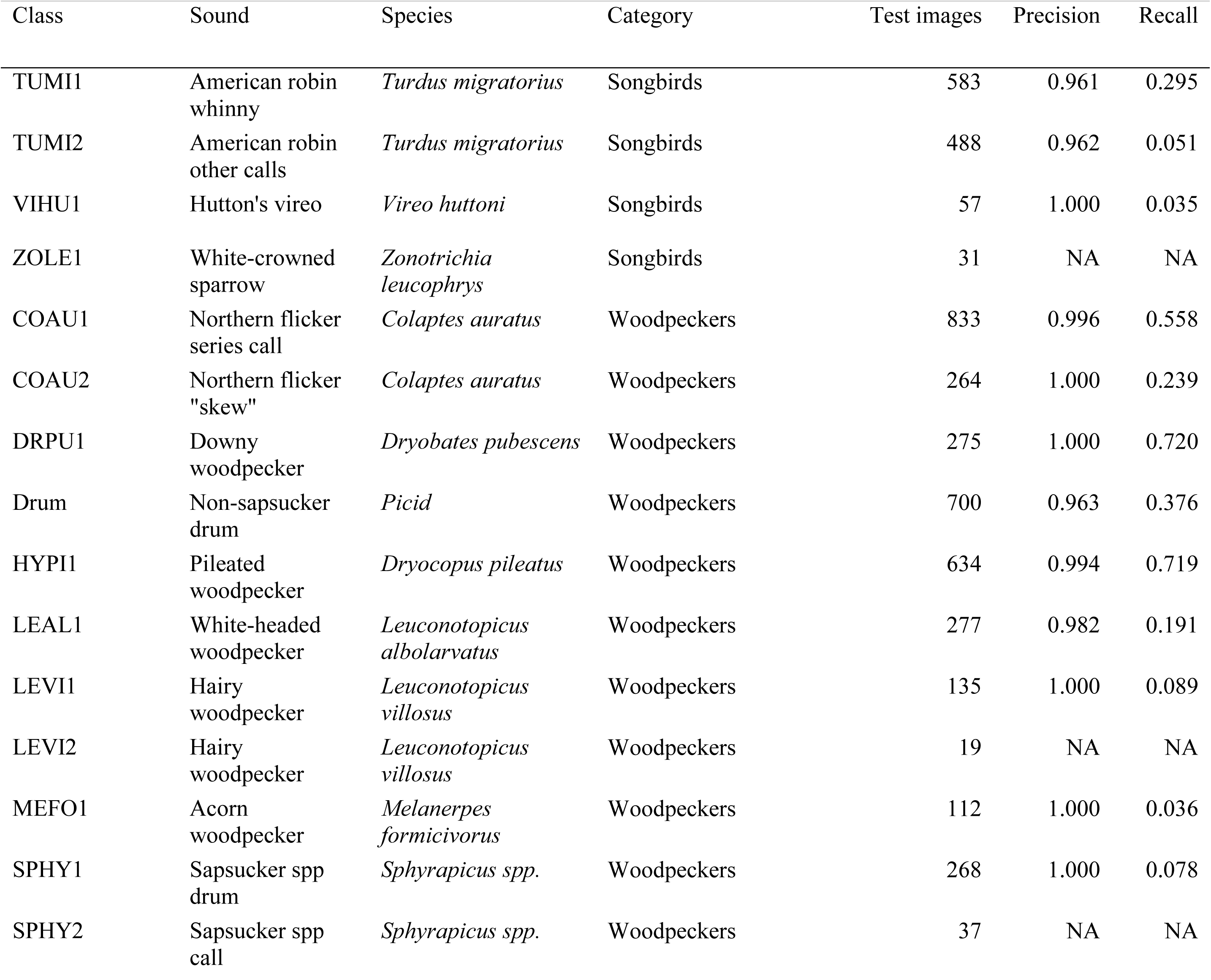
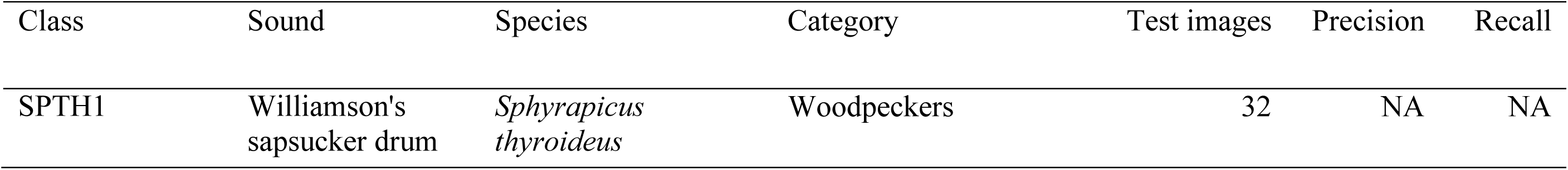
Precision and recall estimates for each sound class of the convolutional neural network (PNW-Cnet v5) used to process bioacoustics data collected during 2023. Unless noted for spotted owl location call, estimates are based on a prediction threshold of 0.95 and were generated with a test set of 41,202 spectrogram images. Estimates for the classes with no apparent detections (images assigned scores ≥ 0.95) in the test set or in the 2023 data are denoted as NA.

### Focal species detections

In 2023, we confirmed ≥1 northern spotted owl detection in all 20% sampling areas and in each of the 2% samples within each state. We consistently found many more male than female detections (Table 5). California study areas (MAR, NWC) had the highest proportion of hexagons with detections (Table 6). Areas in the Washington Cascades (CLE, RAI, WA 2%), Tyee (TYE), and the Oregon Coast Range (COA) had some of the lowest proportion of hexagons with detections (Table 6). The proportion of hexagons with detections has remained consistent on most study areas with multiple years of surveys. TYE is the exception with proportion declining from 0.38 to 0.15 from 2021 to 2023. We did not detect northern spotted owl in Deschutes National Forest, or North Cascades National Park. Indeed, we also did not detect any northern spotted owl north of US Highway 2 in the Washington Cascades (39 hexagons).

**Table 5.**
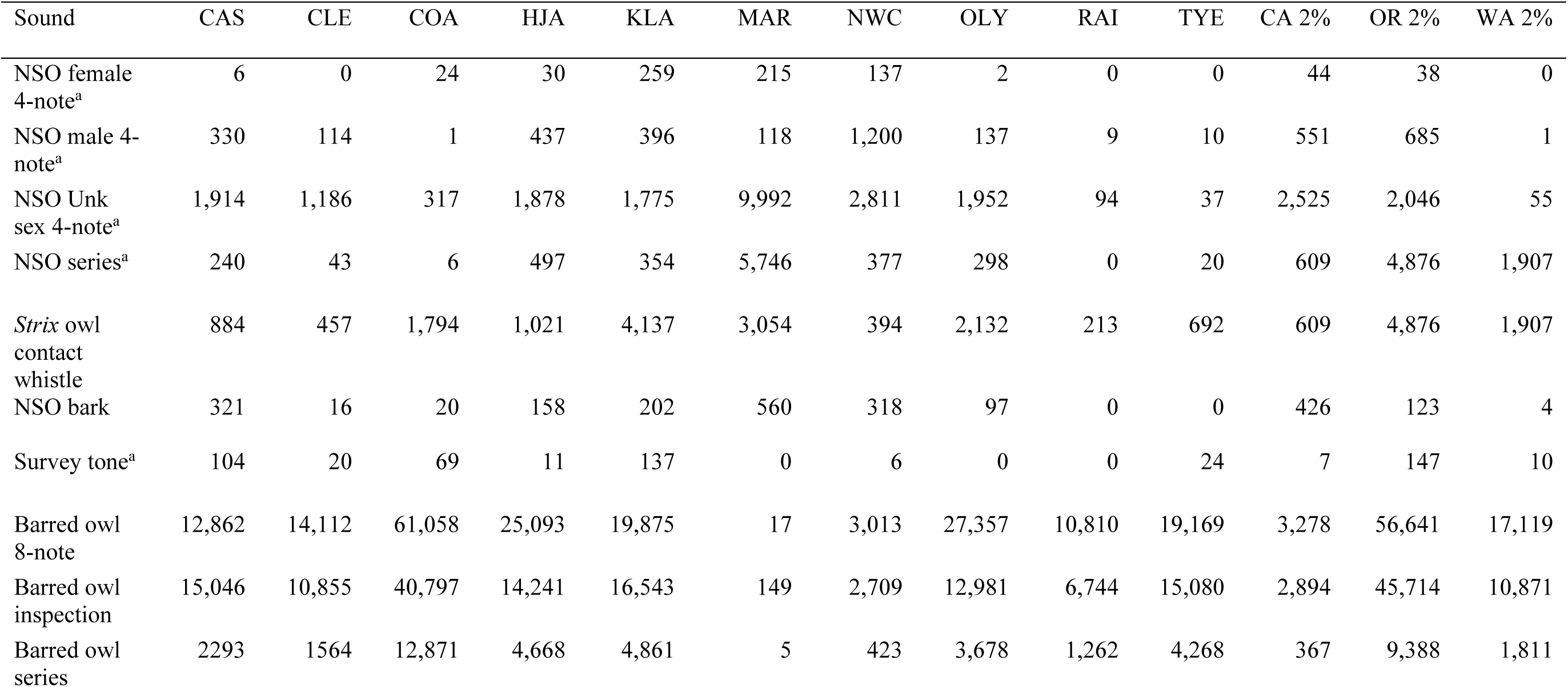
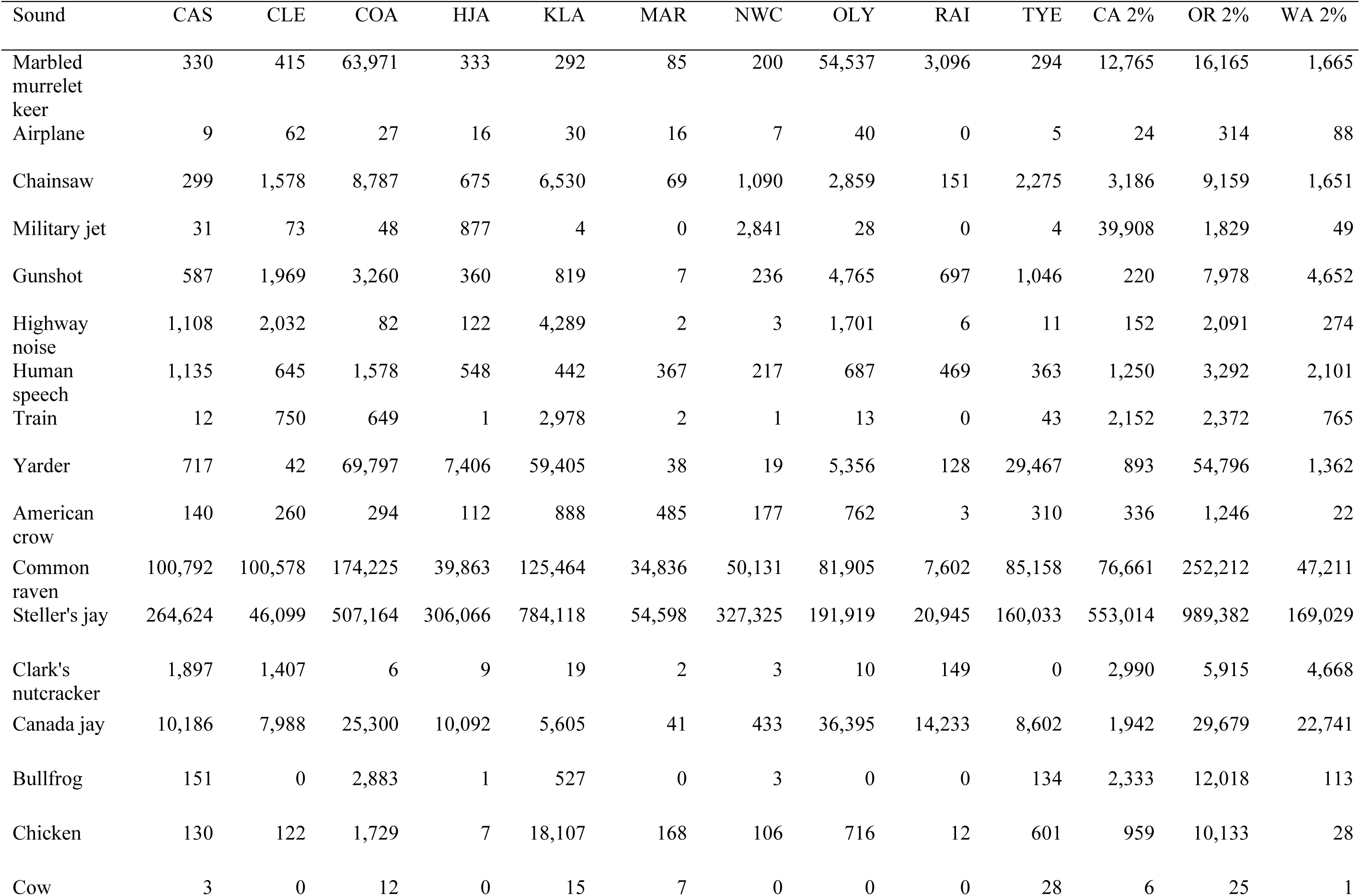
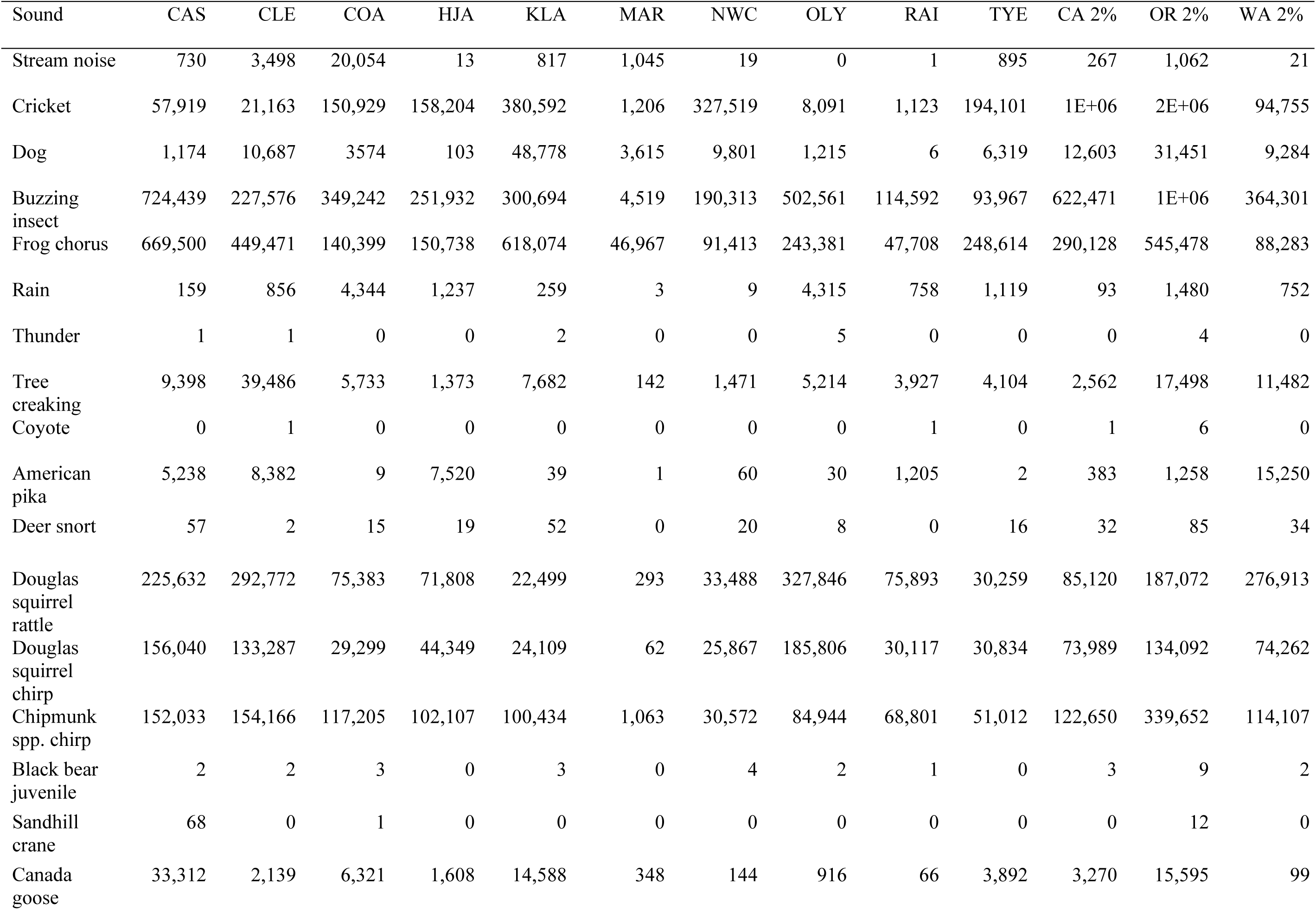
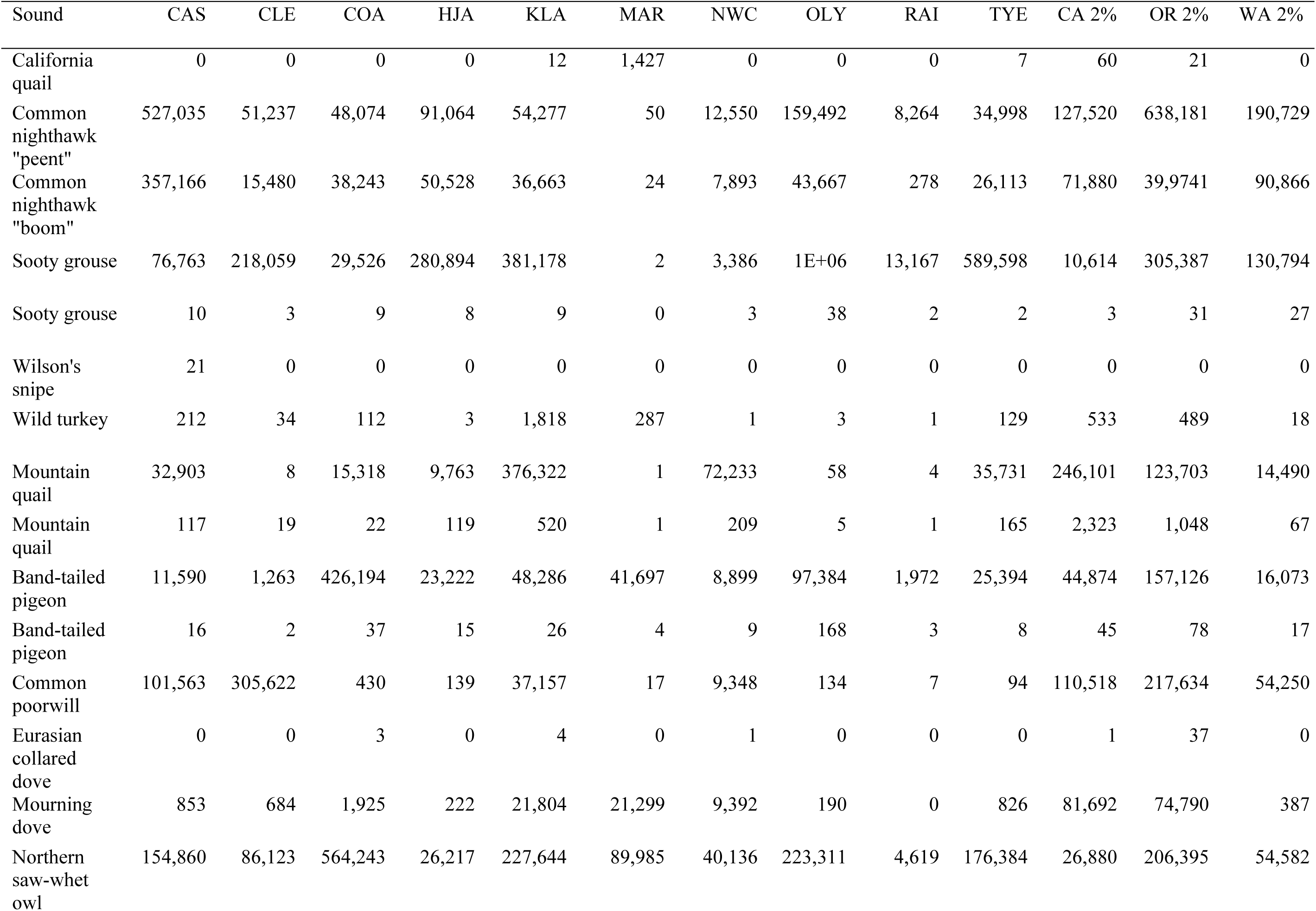
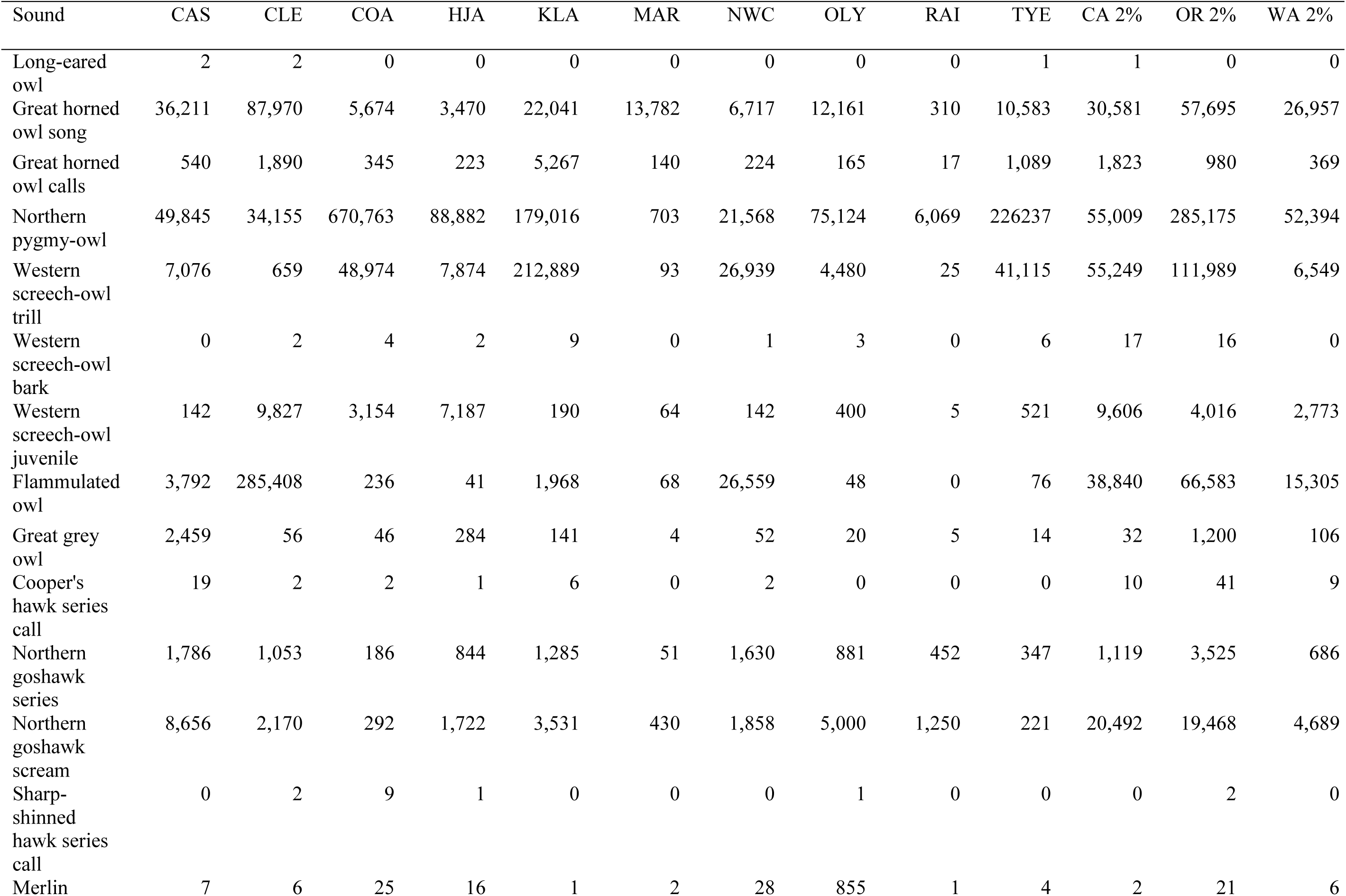
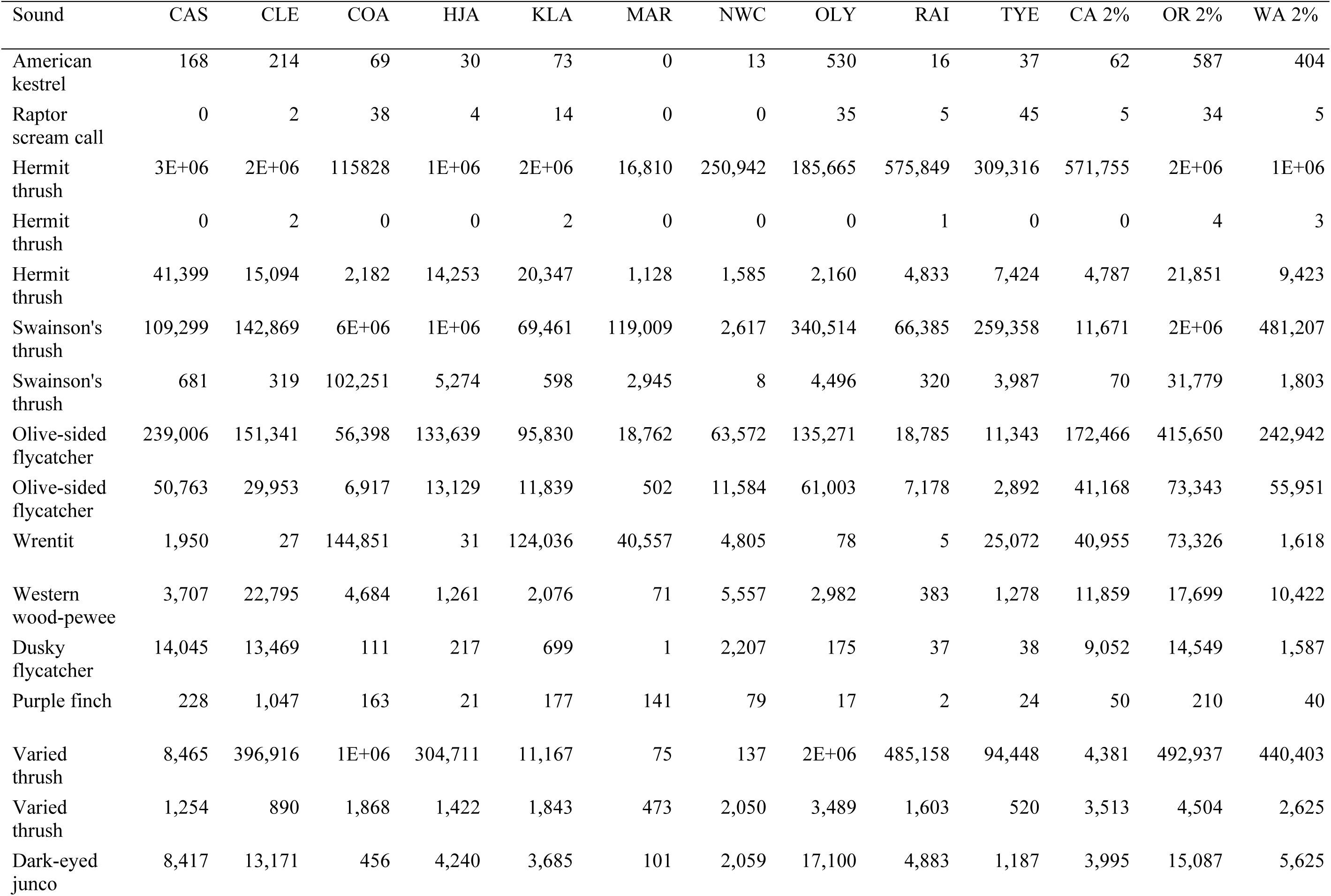
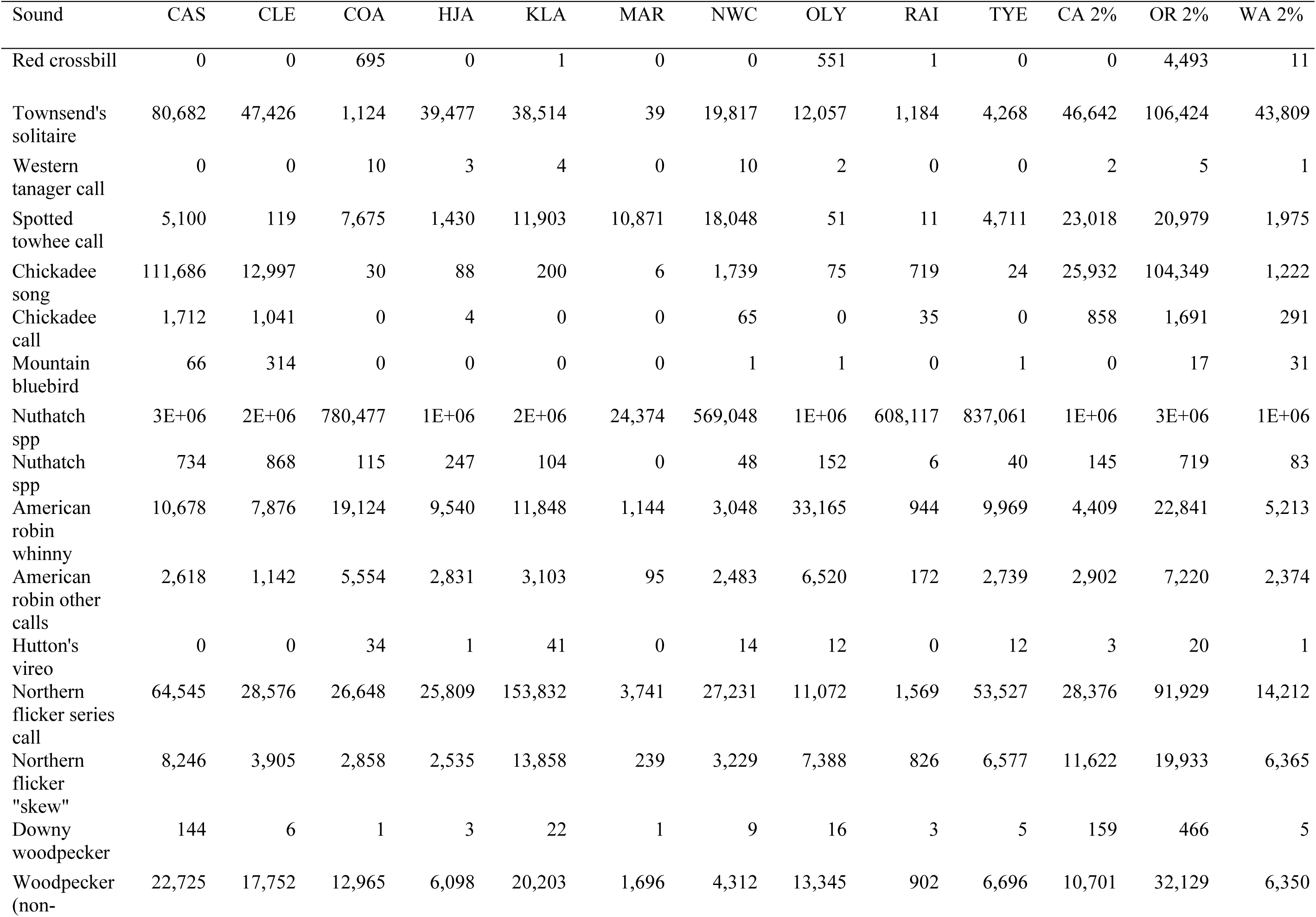
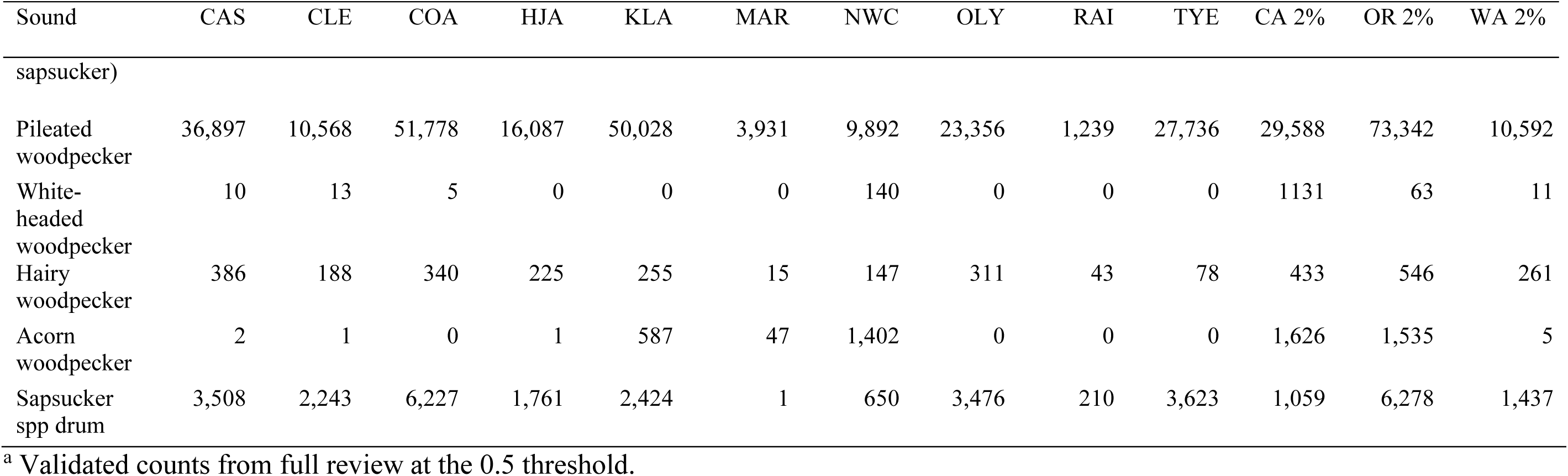
Estimated number of detections of PNW-Cnet v5 sound classes from passive acoustic monitoring data by study area collected in 2023. Estimated detections for each sound class were calculated as the number of 12 s clips in the audio dataset to which the PNW-Cnet v5 assigned a score exceeding 0.95 for that class, multiplied by the precision estimate (Table 4). Validated call counts are given for the primary northern spotted owl (NSO) and survey tone call classes which were fully reviewed at the 0.5 threshold. CAS = Southwest Cascades, OR; CLE = Cle Elum, WA; COA = Coast Range, OR; HJA = H.J. Andrews Experimental Forest, OR; KLA = Klamath Range, OR; NWC = Northwest California, CA; OLY = Olympic Peninsula, WA; RAI = Mount Rainier, WA; TYE = Tyee, OR; WA 2%= 2% random sample of federal lands in Washington; OR 2 = 2% random sample of federal lands in Oregon; CA 2= 2% random sample of federal lands in California. See Table 4 for species and class codes.

**Table 6.** Proportion of monitored hexagons with validated detections of northern spotted owl for years that surveys were conducted (2018–2023) within the Northwest Forest Plan Area.

| Area | 2018 | 2019 | 2020 | 2021 | 2022 | 2023 |
| --- | --- | --- | --- | --- | --- | --- |
| COA | 0.16 | 0.13 | 0.09 | 0.15 | 0.09 | 0.11 |
| OLY | 0.16 | 0.17 | 0.24 | 0.19 | 0.27 | 0.25 |
| KLA |  | 0.43 | 0.53 | 0.49 | 0.51 | 0.37 |
| CLE |  |  | 0.19 | 0.19 | 0.11 | 0.20 |
| CAS |  |  |  | 0.35 | 0.31 | 0.23 |
| HJA |  |  |  | 0.39 | 0.39 | 0.43 |
| MAR |  |  |  | 1.00 | 1.00 | 1.00 |
| NWC |  |  |  | 0.77 | 0.73 | 0.80 |
| RAI |  |  |  | 0.09 | 0.04 | 0.06 |
| TYE |  |  |  | 0.38 | 0.28 | 0.15 |
| WA 2% |  |  |  |  | 0.00<sup>a</sup> | 0.09 |
| OR 2% |  |  |  |  | 0.54<sup>a</sup> | 0.24 |
| CA 2% |  |  |  |  |  | 0.37 |

<sup>a</sup> In 2022, Gifford Pinchot National Forest in Washington and Umpqua National Forest in Oregon were sampled at 2%.

Northern spotted owl call-broadcast surveys were reported, and we detected the survey tone throughout the study region.

Marbled murrelet keer calls were confirmed in COA, OLY, KLA, RAI, TYE, and 2% areas within the range of murrelet in 2023. Marbled murrelets were commonly detected (91-92% of hexagons) on the two most coastal study areas (COA, OLY) (Table 7). A higher proportion of the Oregon 2% sampled areas had murrelet detections compared to Washington and California (Table 7).

**Table 7.** Proportion of monitoring hexagons with validated detections of marbled murrelet for years that surveys were conducted (2018–2023) within the Northwest Forest Plan Area.

| Area | 2018 | 2019 | 2020 | 2021 | 2022 | 2023 |
| --- | --- | --- | --- | --- | --- | --- |
| COA | 0.83 | 0.75 | 0.79 | 0.82 | 0.86 | 0.91 |
| OLY | 0.78 | 0.85 | 0.84 | 0.87 | 0.85 | 0.92 |
| KLA |  | 0.00 | 0.00 | 0.01 | 0.00 | 0.01 |
| CLE |  |  | 0.00 | 0.00 | 0.00 | 0.00 |
| CAS |  |  |  | 0.00 | 0.00 | 0.00 |
| HJA |  |  |  | 0.00 | 0.00 | 0.00 |
| MAR |  |  |  | 0.00 | 0.00 | 0.00 |
| NWC |  |  |  | 0.00 | 0.00 | 0.00 |
| RAI |  |  |  | 0.09 | 0.13 | 0.13 |
| TYE |  |  |  | 0.15 | 0.10 | 0.05 |
| WA 2% |  |  |  |  | 0.00<sup>a</sup> | 0.13 |
| OR 2% |  |  |  |  | 0.08<sup>a</sup> | 0.20 |
| CA 2% |  |  |  |  |  | 0.01 |

<sup>a</sup> In 2022, only Gifford Pinchot National Forest in Washington and Umpqua National Forest in Oregon were sampled at 2%.

All study areas had high proportions of hexagons with barred owl detections, except MAR and California 2% areas (Table 8). In Washington, we detected barred owls in >90% of surveyed hexagons on the west side of the Cascade Mountains and in CLE on the east side we detected barred owls at 80% of hexagons (Table 8). In the Oregon 20% study areas, we detected barred owls at over 95% of hexagons (Table 8). The greatest amount of barred owl 8-note calls was recorded in COA and the Oregon 2% areas (Table 5). California sites had consistently lower proportion of hexagons and number of barred owl 8-note calls compared to sites in Oregon and Washington (Table 5, Table 8). Mendocino National Forest was the only surveyed federal management unit with no barred owl calls detected. Barred owl call-broadcast surveys occurred in some California sampling areas.

**Table 8.**
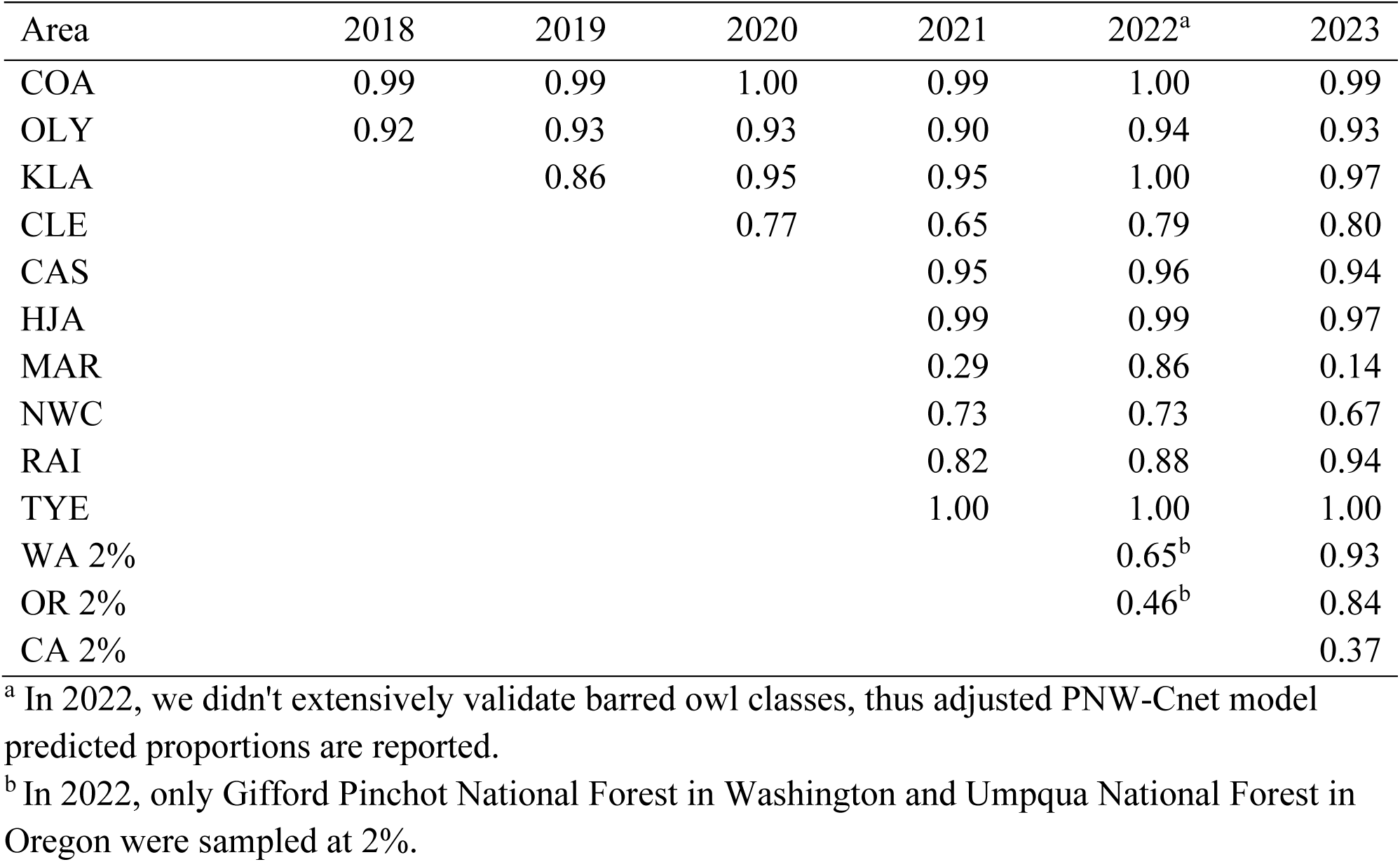
Proportion of monitoring hexagons with validated detections of barred owls for years that surveys were conducted (2018–2023) within the Northwest Forest Plan Area.

Northern saw-whet owls (*Aegolius acadicus*) and northern pygmy owls (*Glaucidium californicum)* had the greatest number of predicted vocalizations throughout our study area (Table 5). We found >1,000 predicted detections of great gray owls in CAS and the OR 2% area (Table 5). Flammulated owls (*Psiloscops flammeolus*) were most commonly detect in CLE, followed by the 2% sample areas in all three states and NWC (Table 5). Great-horned owl (*Bubo virginianus*) and western screech-owl (*Megascops kennicottii*) calls were predicted in all sampling areas (range: 310–87,970 and 25–212,889, respectively). We found a few potential long-eared owl (*Asio otus*) calls in CAS, CLE, TYE, and CA and OR 2% areas (Table 5).

Northern goshawk (*Accipiter gentilis*), a new sound class in PNW-Cnet v5, were classified at the 0.95 model threshold in all areas, with >8,000 classified in CAS, and the CA and OR 2% areas. Pika (*Ochotona princeps*) were commonly (>5,000) predicted in CAS, CLE, HJA, and WA 2%. Band-tailed pigeon (*Patagioenas fasciata*) call classes were found in all areas with as few as 1,263 in CLE up to 426,194 naïve detections in COA (Table 5). Sooty grouse (*Dendragapus fuliginosus*) call classes were predicted most densely at OLY (1,030,627) and TYE (589,598).

We did not confirm detections for wolves (*Canus lupis*), elk (*Cervus canadensis*), ruffed grouse (Bonasa umbellus), red-tailed hawks (*Buteo jamaicensis*), bald eagles (*Haliaeetus leucocephalus*), osprey (*Pandion haliaetus*), Wilson’s warblers (*Cardellina pusilla*), evening grosbeaks (*Hesperiphona vespertina*), orange-crowned warblers (*Leiothlypis celata*), black-headed grosbeaks (*Pheucticus melanocephalus*), chipping sparrows (*Spizella passerine*), pine siskins (*Spinus pinus*), house wrens (*Troglodytes aedon*), or white-crowned sparrows (*Zonotrichia leucophrys*). Each of these species are new classes for PNW-Cnet with small training datasets, so the lack of detections may be from poor model performance rather than absence from areas surveyed. For many of these species the recall in PNW-Cnet v5 may have been too low for any true positives to reach the 0.95 prediction threshold.

## 5. Discussion

### Data collection and processing

Here we report data collected and summaries for passive acoustic monitoring since 2018 within the NWFP area. The first year for full implementation of passive acoustic monitoring of the 2+20% sampling design occurred in 2023 (Lesmeister et al. 2021, Lesmeister and Jenkins 2022). Our goal was to survey approximately 1,100 hexagons which would have included all hexagons that were surveyed in 2022 plus all hexagons included in the full 2% sample outside historical study areas. We were able to survey hexagons surveyed in 2022 and most of the expansion hexagons for a total of 1,009 hexagons surveyed and nearly 2.2 million h of recordings processed and detection encounter histories for 66 species.

With many multi-species survey methods there are tradeoffs that must be made in study design between the number of sampling occasions and sites surveyed (Sanderlin et al. 2014). We have demonstrated the ability to resolve many of these tradeoffs by using passive acoustic monitoring with long-duration deployments, random site selection, multiscale clustered sampling, and high-throughput data processing for a wide range of species over large geographic regions. The NWFP passive acoustic monitoring program was designed to ensure effectiveness for tracking trends in northern spotted owl populations (Lesmeister et al. 2021), but by using a random selection of 5-km^2^ hexagons with multiple sampling stations, we have a design suitable for detecting and studying many other forest-adapted wildlife species (Tosa et al. 2021, Lesmeister and Jenkins 2022). For example, Rugg et al. (2023) used data collected in 2021 and PNW-Cnet output to evaluate western screech-owl occupancy and the effect of barred owls.

Using our data from two study areas, Duarte et al. (2024) demonstrated that the combination of passive acoustic monitoring and PNW-Cnet are effective to estimate marbled murrelet intensity of use across broad scales and will significantly enhance long-term population monitoring. Using a subset of our passive acoustic monitoring data, Weldy et al. (2024) annotated dawn chorus recordings for an open-access dataset that is a valuable resource for researchers developing automated identification tools and those studying the relationship of dawn chorus species to the forest environment.

We continue to develop and improve PNW-Cnet as demonstrated by our use of the most recent version to process data collected in 2023. PNW-Cnet v5 is in preparation for publication and will include performance metrics reported here and an easy to install tool that can be used by field biologists to process passive acoustic monitoring data on personal computers. With continued development of PNW-Cnet we anticipate an expansion of its use for many additional applications to address pressing ecological and conservation challenges.

The primary challenges for the 2023 data collection season were field staff hiring constraints, early season (March–April) weather with low-elevation snow, wildfires in August, and general challenges associated with expanding work into areas unfamiliar to field crews. We continue to gain experience with accessing survey sites and seek continual improvement and efficiencies with recently hired permanent fulltime staff in support of the monitoring program. Once the data are collected, the most significant constraint to producing occupancy estimates for northern spotted owl continues to be our ability to identify calls originating from human surveyors using broadcast play-back surveys. Broadcast surveys have been the primary method of determining northern spotted owl occupancy for the last 40 years and are widely used by private, state, tribal, and federal entities. The use of a three-note tone (USFWS 2021) by some northern spotted owl call-back surveyors has greatly enhanced our ability to screen out and identify call-back surveys during data processing with human review and should expedite occupancy estimates for those overlapping survey areas.

In 2023, we expanded the monitoring network through state and federal partnerships. For example, we partnered with Washington Department of Natural Resources and the Mount Baker-Snoqualmie National Forest to conduct passive acoustic monitoring surveys on that forest. We partnered with California Department of Parks and Recreation to design a survey plan and process data from three hexagons on their lands that they surveyed. We continue to actively seek partnerships to expand the monitoring network with no additional cost to the program.

Additionally, data processing tools developed by the program are freely available and we (PNW Bioacoustics Lab) host workshops each fall to train federal biologists to use the tools for project-level surveys. The 30-year NWFP meta-analysis of northern spotted owl populations is in progress and will include the final analyses of demography and the transition to occupancy. It will include 2023 data to produce the first estimates of range-wide occupancy and population for northern spotted owls.

### Northern spotted owl

Generally, the naïve occupancy of northern spotted owl increased from north to south. We found northern spotted owls in 17%, 24%, and 51% of WA, OR, and CA hexagons respectively. We detected no northern spotted owls north of Highway 2 in the Washington Cascades sampling area, where 35 of 39 surveyed hexagons had barred owl calling. This may be evidence of extirpation from a region which has had the longest exposure to barred owls, however, more years of monitoring data will be needed. Encouragingly, we have not observed a dramatic decline in naïve occupancy rates in most study areas where we now have several years of passive acoustic monitoring. The TYE study area is the exception, where we have conducted three years of passive acoustic monitoring and observed a decline in the proportion of hexagons with detections from 0.38 to 0.15 between 2021 and 2023. A detailed analysis is warranted to investigate potential causes of this change in naïve occupancy rate. We used a combination of a linear predictive model (Dale et al. 2022) and expert review to determine sex of northern spotted owl detections. We confirmed 31,326 four-note calls recorded in 2023. Most of those calls were classified as unknown sex (n = 26,582), but in spot review we found that most of the unknown calls are likely males. Confirmed female four-note calls (n = 755) were only 2.4% of all confirmed four-note calls, highlighting that males are far more vocal and, therefore, more detectable than females. This difference in detection of males and females has important implications for interpreting survey results to predict pair status (Appel et al. 2023). A six-week survey of a hexagon that results in only a single confirmed male detection has a 0.47 probability of being a pair but the female was not detected (Appel et al. 2023).

### Marbled murrelet

Naïve occupancy rates of marbled murrelets in the COA and OLY study areas have remained mostly consistent since 2018, and RAI since 2021. The data collected in 2023 provide us with the first broader occupancy rates outside these study areas, but our estimates include areas surveyed outside the expected range of marbled murrelets inland. In future years we will report naïve occupancy rates only for those areas within the expected range.

### Barred owl

Barred owls have been spreading southwards from Canada for the last 40-50 years and are one of the most detected species in our surveys. We observed detections of barred owls in 89% of hexagons in Washington, 93% in Oregon, and 46% of hexagons in California. We detected barred owls in only one of seven hexagons surveyed in MAR study area and in zero of the 13 hexagons surveyed on the Mendocino National Forest. We observed a general pattern of areas with the highest barred owl naïve occupancy rates having the lowest northern spotted owl naïve occupancy rates (Tables 6, 7).

### Biodiversity

We are increasingly confident that the coupling of passive acoustic monitoring with PNW-Cnet provides the foundation for a powerful toolbox for tracking changes and the drivers of northern spotted owl population change. Further, we are now collecting valuable data for many other wildlife species and are working to identify winners and losers in dynamic landscapes (see Rugg et al. 2023). These opportunities would not be possible with a monitoring program based in traditional field methods that are single-species focused (Lesmeister and Jenkins 2022). We have clearly expanded the number of species we can provide useful information to evaluate population trends and landscape associations, but the monitoring design will not be suitable for all species. Therefore, additional study is needed to resolve which additional species or communities can be integrated into the standardized flow of information generated by the monitoring program. The list of species to be included will be limited to the taxa appropriate to the spatial and temporal (i.e., the target species is forest-adapted and the breeding season range overlaps the NWFP area) sampling design and the pool of species that can be included in automated systems.

We also have several human disturbance noise classes in PNW-Cnet v5 (Table 4). These anthrophony noise classes can provide information on landscape context of our sampling areas. For example, the Oregon study areas, COA, KLA, TYE, and the 2% OR landscape have far more naïve logging yarding system whistle (i.e., yarder’ machine) detections (>50,000) compared to other regions (<7,000). These data will be increasingly informative with long-term monitoring to quantify occupancy dynamics in relation to these disturbance indicators. We will likely be able to generate new insights into these ecosystems due to the flexibility and scale of the biodiversity monitoring program.

## 6. Acknowledgments

Funding and support for this program was provided by: USDI Bureau of Land Management *and* National Park Service; USDA Forest Service Pacific Northwest Region *and* Pacific Northwest Research Station. We thank T. Levi, C. Sullivan, J. Koning, B. Padmaraju, M. Samaduroff, and A. Subramanian for assistance in PNW-Cnet development and data processing. We are deeply indebted to the dedicated field and lab technicians that collected and processed millions of hours of bioacoustics:

2018 field and lab crew: D. Culp, C. Cardillo, A. Ingrassia, D. Jacobsma, K. McLaughlin, P. Papajcik, R. Justice, T. Garrido, A. Froelich. Z. Farrand, A. Munes, Z. Ruff.

2019 field and lab crew: L. Duchac, S. Nuss, M. Peterman, T. Young, H. Lambert, S. Dale, T. Munger, J. Runjaic, A.M. Doss, L.S Bartholomew, E.S. Eber, J.R. Garrido, E.M. Guzman, T.J. Kay, P.M. Loafman, M.H. McConnell, P.S. Papaczik.

2020 field and lab crew: A. Thomas, O. Hardy, H. Lambert, K. Nelson, V. Berdecia, H. Hester, E. Janasov, T. Munger, B. Anderson, A. Nino de Rivera, J. Witteman, J. Runjaic, T. Kohler, Z. Farrand, A. Munes, L. Kufta-Christie, K. Wert, N. Rugg, A.M. Doss, E.S. Eber, E.H. Graham, E.M. Guzman, P.M. Loafman, N.M. Starling, M. Ruggiero, and C.J. Urnes.

2021 field and lab crew: H. Lambert, M. Ruggiero, T. Kohler, M. Parker, B. Henson, S. Reffler, A. Miera, T. Munger, N. Rugg, K. Nelson, A. Thomas, J. Witteman, K. Wright, C. McCafferty, A. Henderson, J. Hurd, K. Wert, D. Culp, J. Fry, N. Murphy, E. Sutphin, L. Platt, E. Tevini, U. Briggs, O. Awbrey, H. Hester, L. Kufta-Christie, R. Neil, J. Runjaic, C. Woods, T. Darling, C. Zeller-Edmonds, A. Doss, E. Eber, E. Graham, E. Guzman, P. Loafman, H. Nay, and C. Urnes.

2022 field and lab crew: L. Platt, S. Reffler, M. Nickols, A. Pastuszek, B. Henson, T. Munger, K. Nelson, J. Schmidt, N. Mutchler, B. Begay, M. Van Bemmel, C. McCafferty, M. Linnell, D. Culp, N. Murphy, J. Fry, C. Armstrong, A. Mueller, E. Tevini, H. Lambert, M. Ruggiero, E. Ormand, K. Wert, A. Henderson, S. Campbell, E. Lessig, J. Runjaic, N. Rugg, H. Hester, and R. Neil.

2023 field and lab crew: J. Crawford, H. Hester, A. Habib, C. Gates, C. Stephens, S. Herring, L. Anderson, K. Nelson, J. Schmidt, B. Begay, T. Christopher Handy, M. Henderson, M. Thelen, B. Norbury, C. Hnilica, J. Fisher, G. Ferone, E. Barnett, M. Murr, A. Pastuszek, R. Hart, E. Lessig, R. Pechtimaldjian, P. Soldi, S. Campbell, N. Murphy, K. Ware, N. McClain, L. Thomas, C. Tescher, N. Mutchler, E. Yargeau, N. Baron, S. Diaz, M. Groves, R. Farrell, and C. Provost.

## Notes

### Competing Interest Statement

The authors have declared no competing interest.

